# Thalamocortical architecture shapes structural and functional plasticity after early sensory loss

**DOI:** 10.64898/2026.08.05.743029

**Authors:** Monami Nishio, Xingyu Liu, Yangwen Xu, Maria Zimmermann, Marcin Szwed, Oliver Collignon, Allyson P. Mackey, Michael J. Arcaro

## Abstract

Early sensory loss transforms the functional organization of affected sensory systems, with regions that normally support vision or audition participating in noncanonical perceptual and cognitive functions. Such reorganization requires coordination across distributed networks. Higher-order thalamic nuclei are positioned to support these interactions through their widespread reciprocal cortical connectivity, yet studies of sensory-loss plasticity have focused primarily on cortex and first-order sensory pathways. Here, we combined structural and functional MRI in early blind and early deaf adults to test how reorganization after sensory loss is expressed across thalamocortical networks. In blindness, structural differences were concentrated along the primary input pathway, with reduced lateral geniculate nucleus volume that covaried with areal organization of primary visual cortex. Functional reorganization was more broadly distributed, with visual cortex showing greater similarity with higher-order (association) thalamic nuclei linked to cognitive control, paralleling a shift toward cognitive-control networks across cortex. This pattern was amplified during a nonvisual perceptual judgment task. Across visual cortex, cross-network functional differences were strongest in regions showing the largest structural effects. A related shift toward control-associated higher-order thalamus was observed in deafness, indicating that higher-order thalamic involvement generalizes across sensory modalities. Together, these results indicate that thalamocortical architecture shapes both the developmental consequences of early sensory loss and the capacity for distributed functional reorganization.

**Highlights:**

- Structural reorganization tracks first-order thalamocortical architecture
- Higher-order thalamus participates in cross-network functional plasticity
- Structural and functional thalamocortical reorganization are anchored to the primary cortical target of sensory input
- Control-related thalamocortical plasticity generalizes across blindness and deafness

**eTOC blurb:** Nishio et al. show that early sensory loss differentially reorganizes first-order and higher-order thalamocortical systems. First-order pathways track cortical structural differences, whereas higher-order thalamus shows cross-network functional reorganization that is enhanced during nonvisual tasks in blindness. Control-related thalamocortical reorganization generalizes to deafness.

## Introduction

Early sensory loss provides a powerful model for studying how experience shapes brain organization. Early blindness and deafness alter the development of sensory pathways and change how affected cortices participate in perception and cognition^1,2^. In blindness, occipital visual cortex is recruited during tactile, auditory, language, and memory tasks^3–9^, while in deafness, auditory cortex is recruited during visual and somatosensory processing^9–16^. These changes are accompanied by altered functional relationships with distributed perceptual and cognitive networks^17–20^. Early sensory loss therefore not only reorganizes processing within the affected sensory system but requires coordination across systems that are only weakly coupled during typical sensory processing.

Higher-order thalamic nuclei are positioned to support this coordination through their widespread reciprocal cortical connectivity with association cortex^21^. The pulvinar interacts broadly with visual, temporal, parietal, cingulate, insular, and prefrontal regions^22–24^, while the mediodorsal nucleus and related association nuclei link the thalamus with prefrontal and frontoparietal systems involved in higher cognition^25–28^. In early blindness, language-sensitive occipital regions show stronger resting-state connectivity with frontal language regions and with thalamic regions identified as ventral lateral and mediodorsal (MD) nuclei^6^. These observations suggest that cross-network plasticity may extend across cortex and higher-order thalamus, but this aspect of sensory-loss reorganization has received far less attention than cortical plasticity itself.

Sensory-system architecture may also constrain where and how this functional reorganization is expressed. First-order, or primary, thalamic nuclei provide the major ascending input to primary sensory cortex and contribute to cortical specification, expansion, and maturation during development^29,30^. Early removal of retinal input in non-human primates alters visual thalamic nuclei, thalamocortical projections, and cortical specification^31–35^. Comparable subcortical changes occur in other sensory systems and at other developmental stages. For example, perinatal deafness in felines alters the volume of subcortical auditory nuclei^36^, and adult somatosensory deafferentation produces delayed reorganization within thalamic and brainstem circuits alongside changes in the primary cortical somatotopic map^37,38^. Human imaging studies likewise show structural differences throughout the primary visual input pathway, from the optic nerves to the lateral geniculate nucleus (LGN) and primary visual cortex (V1), in early blindness^39–43^, and along auditory subcortical pathways in early deafness, although evidence for medial geniculate nucleus reorganization is mixed^36,44,45^. These findings indicate that ascending sensory pathways strongly constrain how sensory systems develop and reorganize in the absence of typical sensory input.

These two aspects of sensory-loss plasticity need not be independent. Following fetal enucleation in macaques, higher-order thalamic projections to visual cortex expand as LGN input is reduced^46^, suggesting that altered sensory input can reshape the balance between ascending and distributed thalamocortical influences. We therefore asked whether early sensory loss reveals complementary constraints and capacities within the same broad architecture. We predicted that structural differences would be most closely related to first-order sensory pathways and the organization of primary sensory cortex, whereas cross-network functional differences would extend into higher-order thalamic systems. We further asked whether these cross-network functional patterns become more pronounced when affected sensory regions are engaged during nonvisual cognition and whether their expression generalizes across sensory modalities.

Here, we combined whole-brain structural and functional MRI in early blindness and early deafness to characterize how sensory loss alters organization across thalamocortical systems. In blindness, we first characterized structural differences along the primary visual pathway and related first-order visual thalamic structure to multiple dimensions of V1 morphology. We then tested whether cross-network functional reorganization extends across cortical and higher-order thalamic networks, and whether structural and functional differences share a common spatial organization across visual cortex. We next asked whether this functional reorganization is modulated during active nonvisual cognition. Finally, we applied parallel structural and functional analyses in early deafness to determine which features of this organization are shared across sensory losses and which depend on the architecture of the affected sensory system.

## Results

### Structural reorganization in blindness is concentrated along the primary visual pathway

We first asked how early blindness alters the structural organization of the primary visual input pathway and whether thalamic and cortical structural differences are related across individuals. We analyzed structural MRI data from 15 early blind participants and 19 neurotypical controls. To localize structural differences without restricting the analysis to predefined regions, we computed whole-brain voxel-wise Jacobian deformation fields^47^ derived from nonlinear registration of individual T1-weighted images to MNI template. Because individual anatomy was warped to the template, positive log-Jacobian values indicate local expansion during registration and therefore smaller native anatomy relative to the template. Negative values indicate local compression and therefore larger native anatomy. A value of zero indicates no local volumetric deformation. Within the thalamus, these template-space deformation effects were summarized across nuclei defined using the THalamus Optimized Multi-Atlas Segmentation (THOMAS) atlas^48^. As a complementary analysis, we also used THOMAS to segment thalamic nuclei in each participant’s native anatomical space and directly estimate nucleus volumes.

Within the thalamus, blind participants showed significantly higher log-Jacobian deformation values (greater local expansion from registration to reference MNI), localized to the lateral geniculate nucleus (LGN; 1,000-permutation test, *P* < 0.001; Fig. 1A). The THOMAS-based nucleus-level analysis confirmed that blind participants had higher LGN deformation values than neurotypical participants (*t*(32) = 4.233, *d = 1.46, P* < 0.001, *P*_FDR_ = 0.002; Fig. 1B), consistent with previous reports of reduced LGN volume in early blindness^39–42,49–53^. No other thalamic nucleus showed a significant group difference in deformation values, including the pulvinar, a higher-order thalamic nucleus extensively connected with visual cortex (*t*(32) = 1.192, *d* = 0.41, *P* = 0.242, *P*_FDR_ = 0.708; Fig. 1B). As a confirmatory analysis, we used THOMAS-based segmentations to directly estimate the volume of each thalamic nucleus in individual space, normalized to the mean volume of that nucleus in the neurotypical group. This analysis similarly showed reduced LGN volume in blind participants, with no detectable volume differences in other nuclei, including the pulvinar (LGN *t*(32) = −4.196, *d* = −1.45, *P* < 0.001, *P*_FDR_ = 0.002; Fig. S1A). No group differences were observed after subdividing the pulvinar into ventral, dorsal, and dorsomedial regions based on prior functional mapping in sighted adults^54^ (v-Pul: *t*(32) = −1.638, *d* = −0.57, *P* = 0.111; d-Pul: *t*(32) = −1.140, *d* = −0.39, *P* = 0.263; dm-Pul: *t*(32) = - 0.916, *d* = −0.32, *P* = 0.367; Fig. S1B, C). In addition to the LGN, we observed significant group differences in Jacobian deformation localized along the optic tract connecting the eyes to the LGN, as well as along the optic radiations connecting the LGN to the primary visual cortex (Fig. S2). Thus, thalamic structural differences between blind and neurotypical participants were focal to the primary thalamic input to cortex.

**Figure 1.**
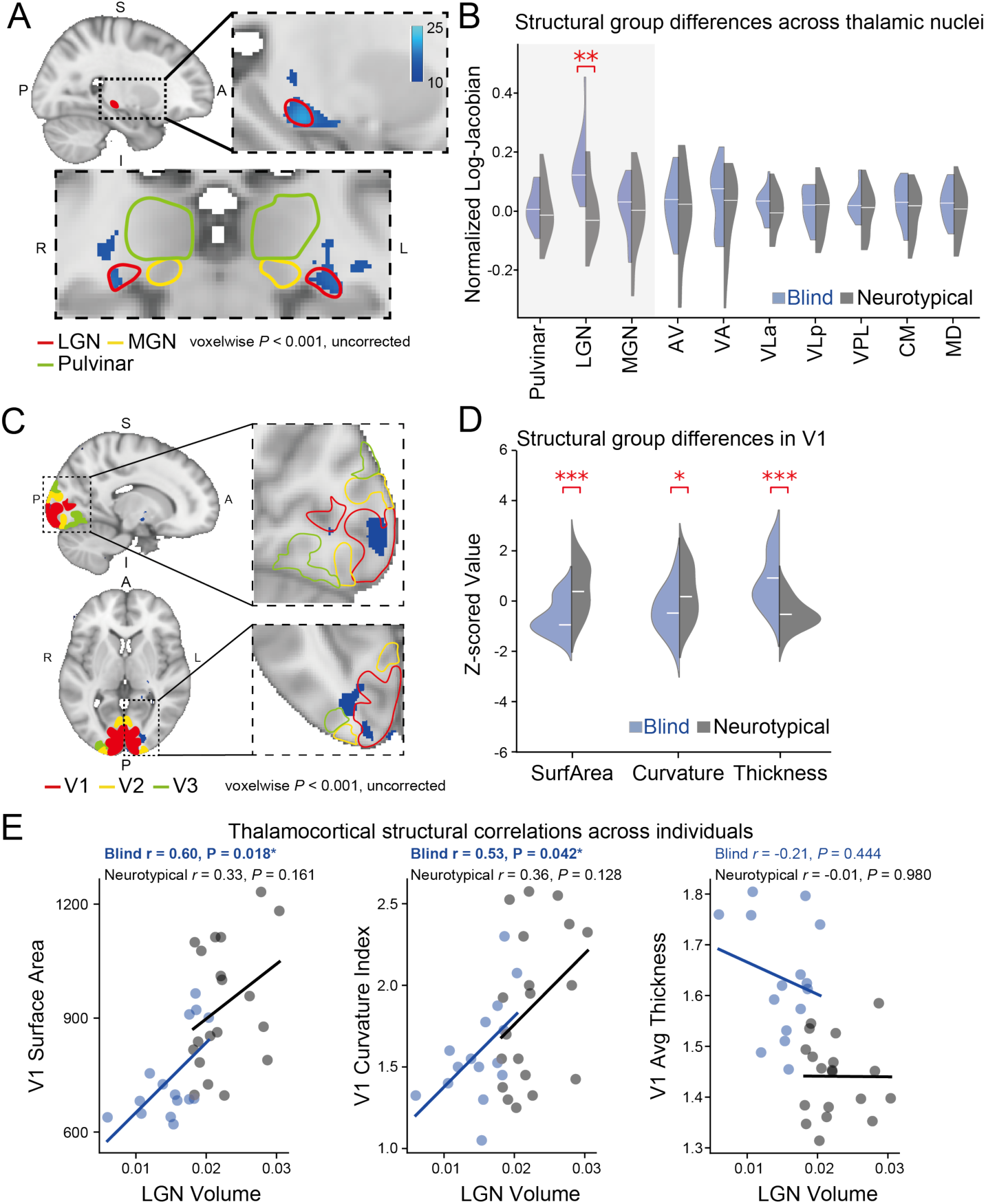
Thalamic and cortical structural reorganization and its relationships in early blindness. **(A)** Voxels highlighted in blue showing significantly higher Jacobian values in blind compared with neurotypical participants were localized to the LGN, with no significant effects in the neighboring medial geniculate nucleus (MGN) or pulvinar (1,000 permutation tests, *P* < 0.001). Log-Jacobian values were derived from the deformation required to register individual anatomy to the MNI template. Higher values indicate greater local expansion to the template and therefore smaller native anatomy relative to the template. **(B)** Participant-level mean log-Jacobian values within each thalamic nucleus in blind and neurotypical participants, mean-centered by subtracting the neurotypical group mean for each nucleus. Zero corresponds to the neurotypical group mean, whereas white lines indicate group medians and therefore need not fall at zero. Visual- and auditory-related nuclei are highlighted in gray (*P* *** < 0.001, ** < 0.01, * < 0.05, FDR-corrected across 10 thalamic nuclei). **(C)** Cortical regions showing significantly higher Jacobian values between blind and neurotypical groups (1,000 permutation tests, *P* < 0.001, uncorrected), highlighted in blue. **(D)** Structural markers in the primary visual cortex (V1) of blind and neurotypical subjects (*P* *** < 0.001, ** < 0.01, * < 0.05, FDR-corrected). FDR correction was applied across the three cortical morphometric features. **(E)** Correlations between LGN volume and structural markers of V1 in blind (blue) and neurotypical (black) participants.

Blindness was also associated with structural differences in visual cortex using the same Jacobian deformation-based approach. The voxel-wise analysis revealed significantly higher deformation values in the blind group than in the neurotypical group, localized to V1 relative to neighboring V2 and V3 as defined by the Wang probabilistic retinotopic atlas^55^ (1,000-permutation test *P* < 0.001; Fig. 1C). We then quantified structural features within V1. Blind participants showed reduced surface area, curvature index (a scale-independent measure of the total intrinsic curvature of a cortical region)^56^, and increased cortical thickness relative to neurotypical participants (surface area: *t*(32) = −3.898, *d* = −1.35, *P* < 0.001, *P*_FDR_ < 0.001; curvature: *t*(32) = −2.083, *d* = −0.72, *P* = 0.045, *P*_FDR_ = 0.045; thickness: *t*(32) = 4.685, *d* = 1.62, *P* < 0.001, *P*_FDR_ < 0.001; Fig. 1D).

We then asked whether individual differences in LGN volume were related to V1 morphometry. In blind participants, LGN volume showed nominal positive associations with V1 surface area and curvature, but not cortical thickness (surface area: *r*(13) = 0.60, *P* = 0.018, *P*_FDR_ = 0.054 [95% CI 0.13, 0.85]; curvature: *r*(13) = 0.53, *P* = 0.042, *P*_FDR_ = 0.063 [95% CI 0.02, 0.82]; thickness: *r*(13) = −0.21, *P* = 0.444, *P*_FDR_ = 0.444 [95% CI −0.65, 0.34]; Fig. 1E). No corresponding associations were detected in neurotypical participants (surface area: *r*(17) = 0.33, *P* = 0.161 [95% CI −0.15, 0.68]; curvature: *r*(17) = 0.36, *P* = 0.128 [CI −0.11, 0.70]; thickness: *r*(17) = −0.01, *P* = 0.980 [95% CI −0.46, 0.45]; Fig. 1E). Together, these results show that early blindness was associated with structural alterations along the primary visual input pathway to cortex, and further reveal, within blind individuals, a selective relationship between first-order visual thalamic structure and areal features of primary visual cortex, but not cortical thickness.

### Cross-network plasticity extends across cortex and higher-order thalamus

The concentration of structural differences along the primary visual pathway does not explain how deprived visual cortex becomes integrated with noncanonical cortical networks. We therefore asked whether early blindness was also associated with altered functional organization across distributed thalamocortical networks. Using resting-state fMRI, we characterized each cortical parcel and thalamic voxel by its pattern of functional relationships with the rest of the cortex (its functional fingerprint)^54^, and compared the similarity of these fingerprint profiles across cortical and thalamic functional networks defined by the Yeo 7-network parcellation^57^. Relative to pairwise time-series correlations, which depend on individual connections, fingerprint profiles summarize the full functional connectivity pattern of a region and therefore are less affected by noise in any single correlation. This is an advantage for the thalamus, where signal-to-noise is low and varies across nuclei with tissue composition.

At the cortical level, visual parcels showed increased functional fingerprint similarity with control network parcels and reduced fingerprint similarity with somatomotor network parcels in blind compared with neurotypical participants, consistent with previous findings^6,7,58^ (control: *t*(32) = 7.635, *d* = 2.64, *P* < 0.001, *P*_FDR_ < 0.001; somatomotor: *t*(32) = −10.389, *d* = −3.59, *P* < 0.001, *P*_FDR_ < 0.001; Fig. 2A). This reduction in visual–somatomotor similarity persisted when the somatomotor network was partitioned into auditory-related and non-auditory subnetworks (Fig. S3). Notably, functional fingerprint similarity between parcels within the visual network did not differ between groups (visual: *t*(32) = −1.020, *d* = −0.35, *P* = 0.315, *P*_FDR_ = 0.368; Fig. 2A), in line with previous resting-state studies showing preserved within-network correlation structure and retinotopic-like organization in early blindness^59,60^. Although not statistically significant, LGN volume and V1 surface area showed negative associations with visual–control fingerprint similarity in blind participants (LGN: *r*(13) = −0.45, *P* = 0.096 [95% CI −0.78, 0.08]; V1 surface area: *r*(13) = −0.50, *P* = 0.056 [95% CI −0.81, 0.02]; Fig. S4A,B), suggesting that reduced bottom-up thalamic input to visual cortex may be associated with greater cross-network functional reorganization of visual cortex.

**Figure 2.**
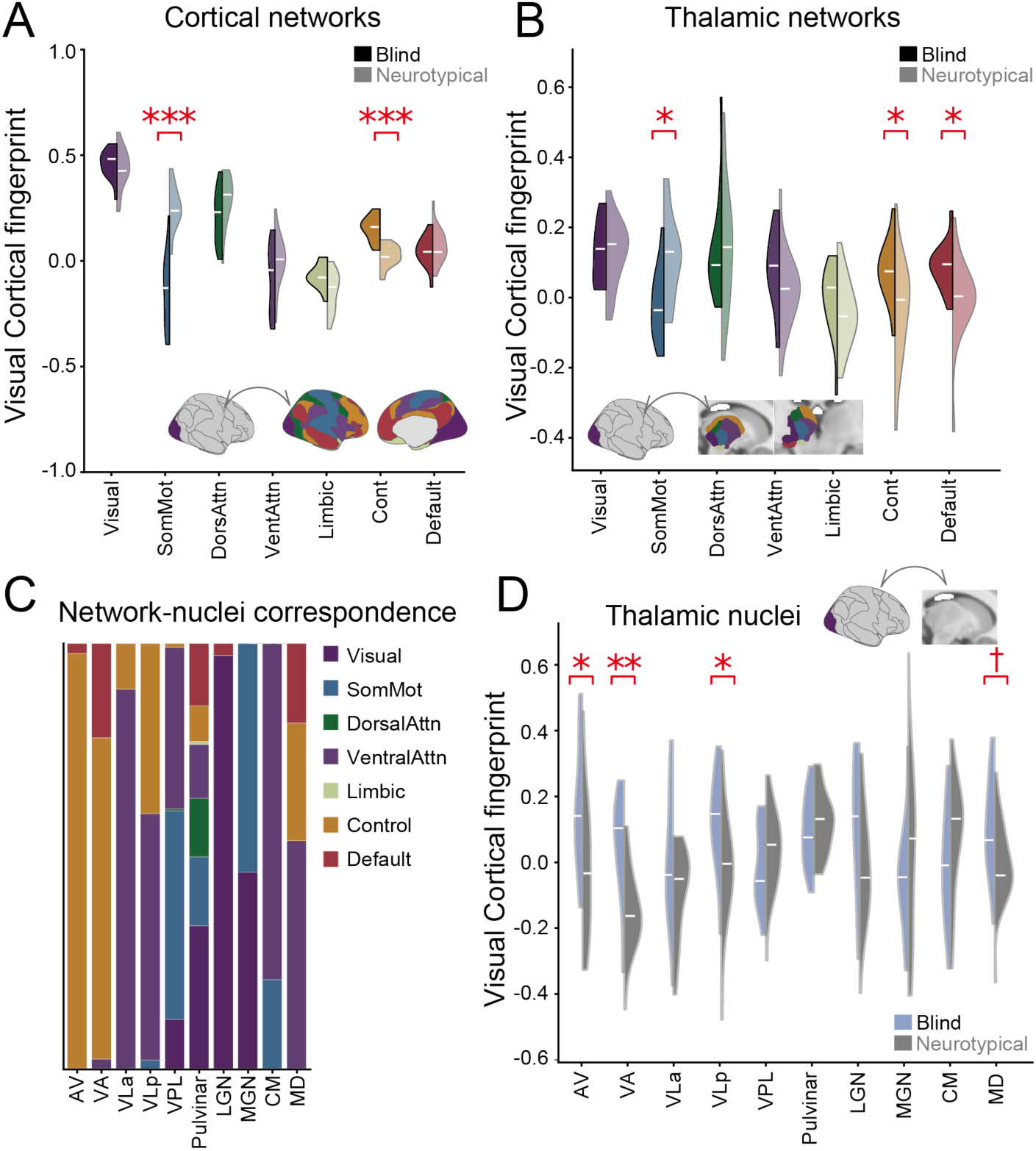
Cortical and thalamic functional reorganization in early blindness. **(A, B)** Functional fingerprint similarity of the cortical visual network with cortical (A) and thalamic (B) Yeo networks in blind and neurotypical groups. \*\*\**P* < 0.001, \*\**P* < 0.01, \**P* < 0.05, FDR-corrected. FDR correction was applied across 7 networks. Cortical and thalamic network distributions are shown in the bottom of each panel. **(C)** Correspondence between Yeo functional networks and thalamic nuclei. **(D)** Functional fingerprint similarity between the cortical visual network and thalamic nuclei. \*\**P* < 0.01 (FDR-corrected), \**P* < 0.05 (FDR-corrected), †*P* < 0.05 (FDR-uncorrected). FDR correction was applied across 10 thalamic nuclei.

We next asked whether this cross-network shift was mirrored in the thalamus. The cortical visual network showed increased fingerprint similarity with thalamic control and default mode networks and reduced similarity with the thalamic somatomotor network in blind compared to neurotypical participants (control: *t*(32) = 2.540, *d* = 0.88, *P* = 0.016, *P*_FDR_ = 0.038; default mode: *t*(32) = 2.673, *d* = 0.92, *P* = 0.012, *P*_FDR_ = 0.038; somatomotor: *t*(32) = −3.311, *d* = −1.14, *P* = 0.002, *P*_FDR_ = 0.016; Fig. 2B). A complementary analysis anchored on thalamic networks showed a similar pattern, with reduced fingerprint similarity between the thalamic somatomotor network and cortical visual parcels and increased fingerprint similarity between the thalamic control network and cortical visual parcels (Fig. S5). Thus, the cross-network pattern distinguishing blind and neurotypical individuals was evident across both cortical and thalamic systems.

At the level of anatomical thalamic nuclei, each nucleus showed a distributed network fingerprint profile (Fig. 2C). Compared with neurotypical participants, visual cortex in blind participants showed increased fingerprint similarity between the cortical visual network and the anterior ventral (AV), ventral anterior (VA), ventral lateral posterior (VLp), and MD nuclei, although the MD effect did not survive FDR correction (AV: *t*(32) = 2.610, *d* = 0.90, *P* = 0.014, *P*_FDR_ = 0.046; VA: *t*(32) = 3.863, *d* = 1.33, *P* < 0.001, *P*_FDR_ = 0.005; VLp: *t*(32) = 2.778, *d* = 0.96, *P* = 0.009, *P*_FDR_ = 0.045; MD: *t*(32) = 2.445, *d* = 0.84, *P* = 0.020, *P*_FDR_ = 0.050; Fig. 2D). Complementary analyses of other cortical networks and thalamic nuclei are shown in Fig. S6. Together, these results show that cross-network functional differences in early blindness are expressed in higher-order thalamic nuclei rather than the first-order sensory input nucleus.

### Structural and cross-network functional differences share a common spatial organization

Because V1 showed both structural relationships with the primary visual thalamus and cross-network functional differences involving higher-order thalamic systems, we asked whether these effects varied systematically across the broader visual cortical hierarchy. Extrastriate visual areas are connected directly and indirectly with V1^61^, and cortical connection probability and strength decline with distance^62,63^. We therefore used geodesic distance from V1 as a spatial measure related to both proximity to the primary cortical target of retinal input and position along the visual cortical hierarchy^64^. We quantified blind-neurotypical differences in surface area, curvature, and cortical thickness across Schaefer cortical parcels^65^ and related these effects to V1 geodesic distance. Cortical parcels were classified as visual or nonvisual based on their overlap with the Wang probabilistic retinotopic atlas^55^, allowing us to test whether this spatial organization was specific to visual cortex.

Across visual parcels, blindness-related structural differences varied systematically with geodesic distance from V1 (Fig. 3A). Reductions in surface area and curvature index were largest near V1 and became smaller with increasing distance (surface area: *r*(84) = 0.60, *P* < 0.001, *P*_spin_ = 0.033; Fig. 3B, curvature: *r*(84) = 0.43, *P* < 0.001, *P*_spin_ = 0.004; Fig. 3C). Cortical thickness showed the complementary pattern, with the largest increases near V1 and smaller differences with increasing distance (*r*(84) = −0.54, *P* < 0.001, *P*_spin_ = 0.008; Fig. 3D). These relationships were not significant after spatial-autocorrelation correction in nonvisual cortex (surface area: *r*(312) = −0.11, *P* = 0.051, *P*_spin_ = 0.546; Fig. 3B, curvature: *r*(312) = 0.04, *P* = 0.455, *P*_spin_ = 0.692; Fig. 3C, thickness: *r*(312) = −0.12, *P* = 0.028, *P*_spin_ = 0.110; Fig. 3D). Structural differences across visual cortex were also related to position along the cortex-wide sensorimotor–association axis (Fig. S7), suggesting that blindness-related structural differences in visual cortex may be situated within a broader cortex-wide hierarchy. Together, these findings indicate that the structural consequences of blindness extend beyond V1 but remain organized with respect to the primary cortical entry point of visual input, consistent with the idea that first-order thalamocortical architecture constrains how visual cortex develops in the absence of visual input.

**Figure 3.**
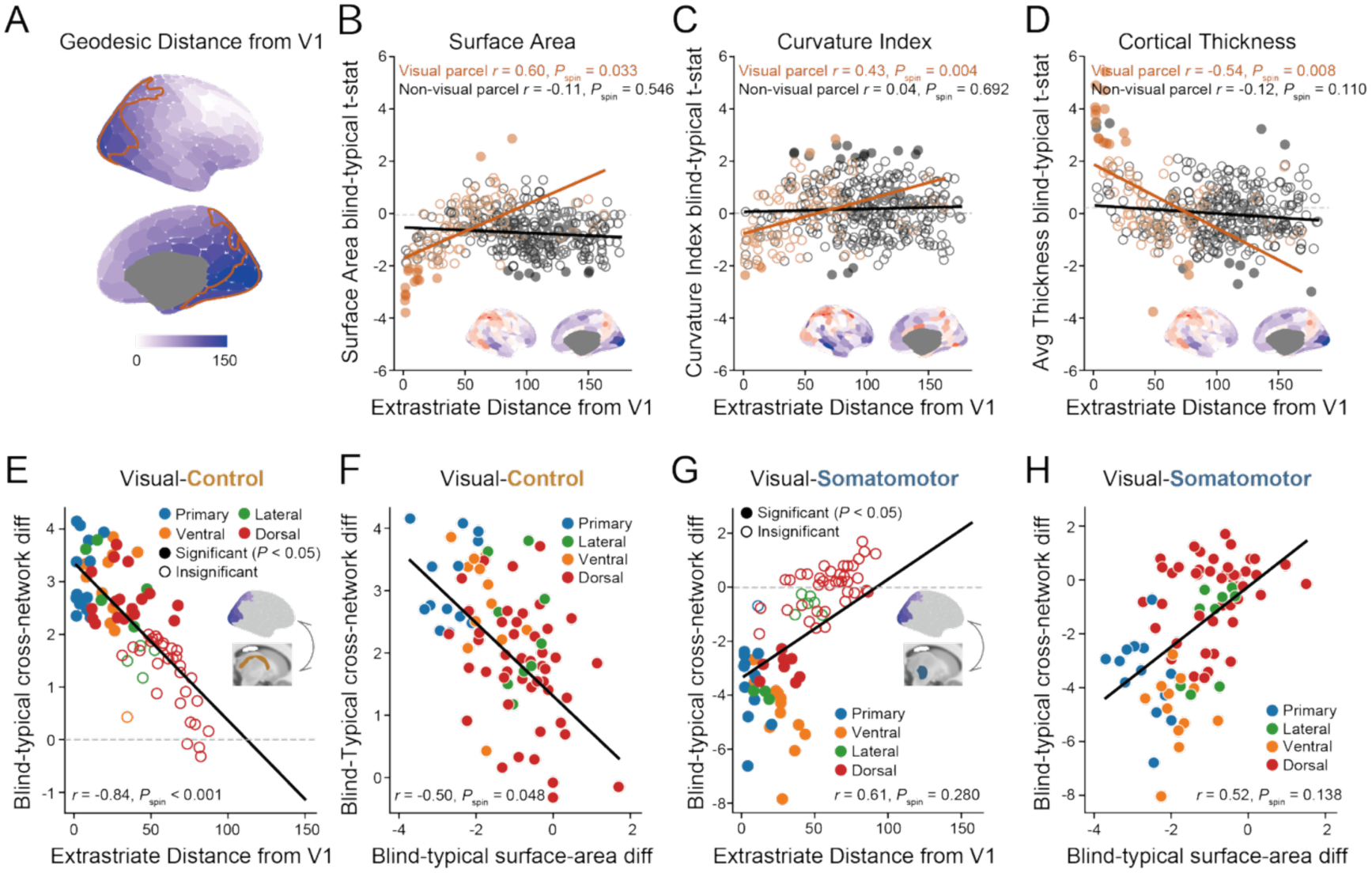
Structural constraints and cross-network plasticity share a V1-anchored cortical organization. **(A)** Geodesic distance from V1, with retinotopic visual cortex outlined in orange. **(B-D)** Blind–neurotypical t-statistics for surface area (B), curvature index (C) and cortical thickness (D) across Schaefer 400 parcels, plotted as a function of geodesic distance from V1. The 86 parcels overlapping visual regions are shown in orange, and non-overlapping parcels in black. Filled dots indicate statistical significance (*P* < 0.05). Surface maps in the bottom-right panels show the spatial distribution of blind–neurotypical *t*-statistics for each feature, standardized to a range of −2 to 2. **(E, G)** Blind–neurotypical t-statistics for visual–thalamic control (E) and visual–thalamic somatomotor (G) functional fingerprint similarity across the same 86 parcels, plotted as a function of geodesic distance from V1. Filled dots indicate statistical significance (*P* < 0.05). Colors denote visual processing streams (blue: primary, orange: ventral, green: lateral, red: dorsal). **(F, H)** Spatial correspondence between structural and cross-network functional differences across visual cortical parcels for visualthalamic control (F) and visual-thalamic somatomotor (H) networks.

We then asked whether the cross-network functional differences also varied systematically with geodesic distance from V1. The increased similarity between visual cortex and thalamic control-network connectivity fingerprints showed a strong spatial gradient, with the largest group differences occurring near early visual cortex (*r*(84) = −0.84, *P* < 0.001, *P*_spin_ < 0.001; Fig. 3E). A similar pattern was observed for the reduced similarity between visual cortex and thalamic somatomotor-network fingerprints (*r*(84) = 0.61, *P* < 0.001), although this association did not survive spatial autocorrelation correction (*P*_spin_ = 0.280; Fig. 3G). Regions showing larger blindness-related reductions in surface area likewise showed greater increases in visual–thalamic control fingerprint similarity (*r*(84) = −0.50, *P* < 0.001, *P*_spin_ = 0.048; Fig. 3F). Visual–thalamic somatomotor differences showed a parallel association with surface area differences (*r*(84) = 0.52, *P* < 0.001), but this relationship did not survive spatial autocorrelation correction (*P*_spin_ = 0.138; Fig. 3H). Finally, visual–thalamic control fingerprint similarity varied significantly across major visual processing streams, with smaller group differences in the dorsal pathway than in ventral and lateral visual cortex (*F*(3,96) = 3.52, *P* = 0.018; Fig. S8). Visual–thalamic somatomotor fingerprint similarity showed a qualitatively similar pattern across streams, but the group × stream interaction did not reach statistical significance (*F*(3,96) = 2.44, *P* = 0.069; Fig. S8). Notably, cortical visual–control and visual–somatomotor fingerprint similarity showed a similar spatial organization across all three analyses, with blindness-related differences varying systematically with V1 geodesic distance, covarying with regional surface area differences, and differing across visual processing pathways (Fig. S9). Together, these findings show that structural constraints and cross-network plasticity share a spatial organization centered on the primary cortical target of thalamic input, with the strongest evidence for enhanced functional associations with the control network. Regions closest to the primary cortical target of first-order visual input showed the strongest structural and cross-network thalamocortical and cortical functional differences, consistent with the idea that the architecture of primary sensory input constrains cortical development while shaping where cross-network plasticity can be expressed.

### Nonvisual task engagement amplifies visual-control relationships in blindness

Given that the resting-state analyses showed stronger fingerprint similarity between visual cortex and control-related regions in blind participants, we next asked whether this effect was further expressed during active nonvisual tasks, when affected visual cortex may contribute to task-relevant perceptual and cognitive processing. We tested this question in the same participants using an auditory object judgment task in which participants judged the shape and function of objects based solely on auditory information, without any visual input (Fig. 4A)^66^. As previously reported^66^, task performance was comparable across groups.

**Figure 4.**
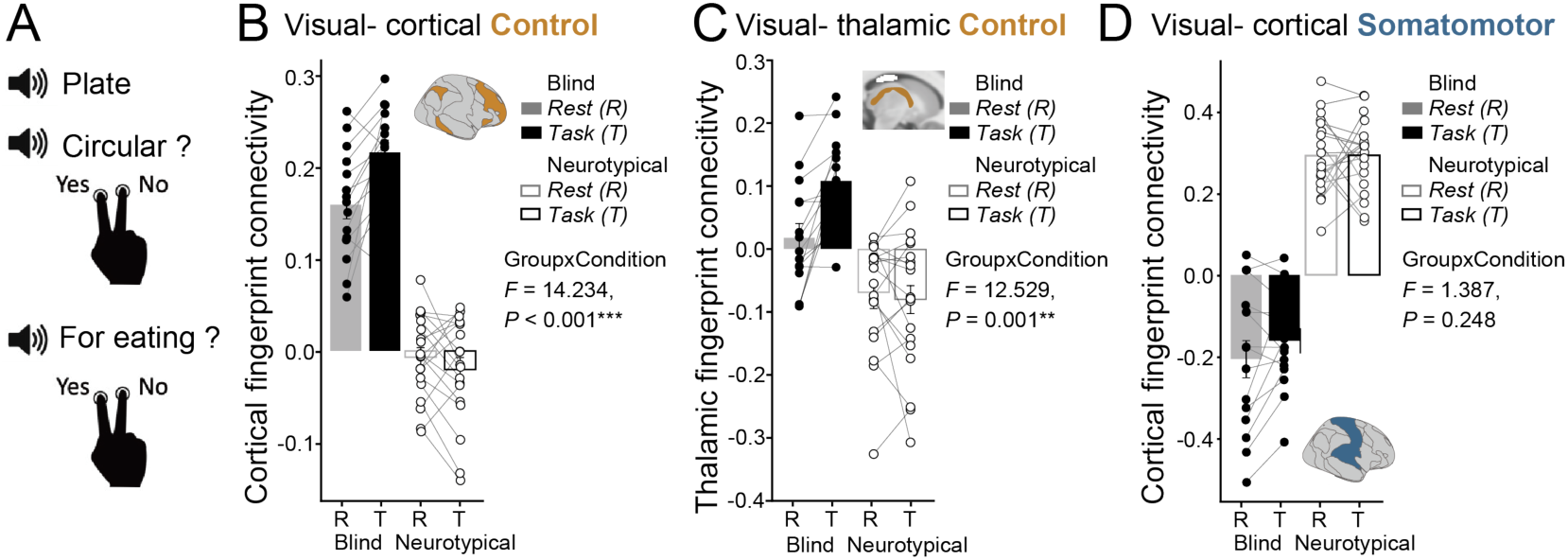
Task-dependent enhancement of visual-control association in cortex and thalamus. **(A)** Auditory object task. Participants judged object shape and function from auditory word stimuli without visual input. **(B)** Visual–cortical control functional fingerprint similarity during rest (R) and task (T) in blind and neurotypical participants. **(C)** Visual-thalamic control functional fingerprint similarity during rest and task in blind and neurotypical participants. **(D)** Visual-cortical somatomotor functional fingerprint similarity during rest (R) and task (T) in blind and neurotypical participants, averaged across Schaefer 400 parcels overlapping visual cortex. Filled circles indicate blind participants, and open circles indicate neurotypical participants.

Visual-control fingerprint similarity showed a significant group x condition interaction (*F*(1, 32) *=* 14.234*, P* < 0.001; Fig. 4B). Relative to the resting-state experiment, visual-control network similarity increased during the task in blind participants, whereas it decreased modestly in neurotypical participants (blind: *t*(14) = −2.756, *dz* = −0.71, *P* = 0.015; neurotypical: *t*(18) = 2.639, *dz* = 0.61, *P* = 0.017; Fig. 4B). Consequently, the blind-neurotypical group difference was greater during the task than at rest (rest: *t*(32) = 7.68, *d* = 2.65, *P* < 0.001; task: *t*(32) = 12.77, *d* = 4.41, *P* < 0.001). At the thalamic level, fingerprint similarity between the cortical visual network and thalamic control network also showed task-related amplification in blind participants (*F*(1, 32) *=* 12.529*, P* = 0.001; Fig. 4C). These effects were specific to the control-related processing, as visual-somatomotor cortical fingerprint similarity showed no significant group x condition interaction (*F*(1, 32) *=* 1.387*, P* = 0.248; Fig. 4D). Together, these results suggest that the visual-control network associations in early blindness are further expressed during active nonvisual cognition at both cortical and thalamic levels.

### Shared higher-order network plasticity across sensory losses

The blindness analyses identified two complementary features of early sensory loss: structural differences associated with the primary sensory input pathway and cross-network plasticity involving higher-order thalamic systems. To test which aspects generalize across modality loss, we next examined thalamocortical organization in early deafness using structural and functional MRI data from 25 early deaf participants and 30 neurotypical controls.

In contrast to blindness, early deafness was not associated with detectable structural differences in any thalamic nucleus, including the MGN, the first-order auditory counterpart of the LGN, in either the voxel-wise or nucleus-level analysis (Fig. 5A,B). The absence of detectable MGN differences may reflect its greater synaptic distance from the cochlea than the LGN is from the retina. Consistent with this interpretation, lower Jacobian values were evident within the right inferior colliculus, which lies one relay closer to the cochlea (Fig. S10). Despite a lack of clear thalamic structural differences, deaf participants nevertheless showed structural differences in primary auditory cortex (TE1: 1,000-permutation test *P* < 0.001; Fig. 5C). As in visual cortex in blindness (Fig. 1C, D), TE1 in deaf participants showed reduced surface area and curvature, along with increased cortical thickness, in TE1 (surface area: *t*(53) = −2.203, *d* = −0.60, *P* = 0.032, *P*_FDR_ = 0.041; curvature: *t*(53) = −2.095, *d* = −0.57, *P* = 0.041, *P*_FDR_ = 0.041; thickness: *t*(53) = 3.985, *d* = 1.08, *P* < 0.001, *P*_FDR_ < 0.001; Fig. 5D). However, the reduction in TE1 surface area and curvature in deaf participants was smaller than the reduction observed in V1 in blind participants (permutation test, surface area: blind-deaf *Diff* = 185, *P* = 0.001, *P*_FDR_ = 0.003; curvature: blind-deaf *Diff =* 0.245*, P* = 0.092, *P*_FDR_ = 0.138), whereas the increase in cortical thickness did not differ between modality-loss groups (blind-deaf *Diff =* 0.028*, P* = 0.724, *P*_FDR_ = 0.724). Reductions in surface area and curvature were strongest near TE1 and decreased with increasing geodesic distance (surface area: *r*(47) = 0.46, *P* = 0.001, *P*_spin_ = 0.039; curvature: *r*(47) = 0.42, *P* = 0.003, *P*_spin_ = 0.007), whereas cortical thickness did not vary significantly with distance (thickness: *r*(47) = −0.11, *P* = 0.464, *P*_spin_ = 0.451; Fig. S11). Unlike the structural pattern in blindness, these effects were not related to the cortex-wide S–A axis (Fig. S11). Compared with blindness, these findings suggest that structural alterations in first-order thalamic nuclei are not a uniform consequence of early sensory loss, whereas an increase in cortical thickness may reflect a more shared feature across sensory-loss groups.

**Figure 5.**
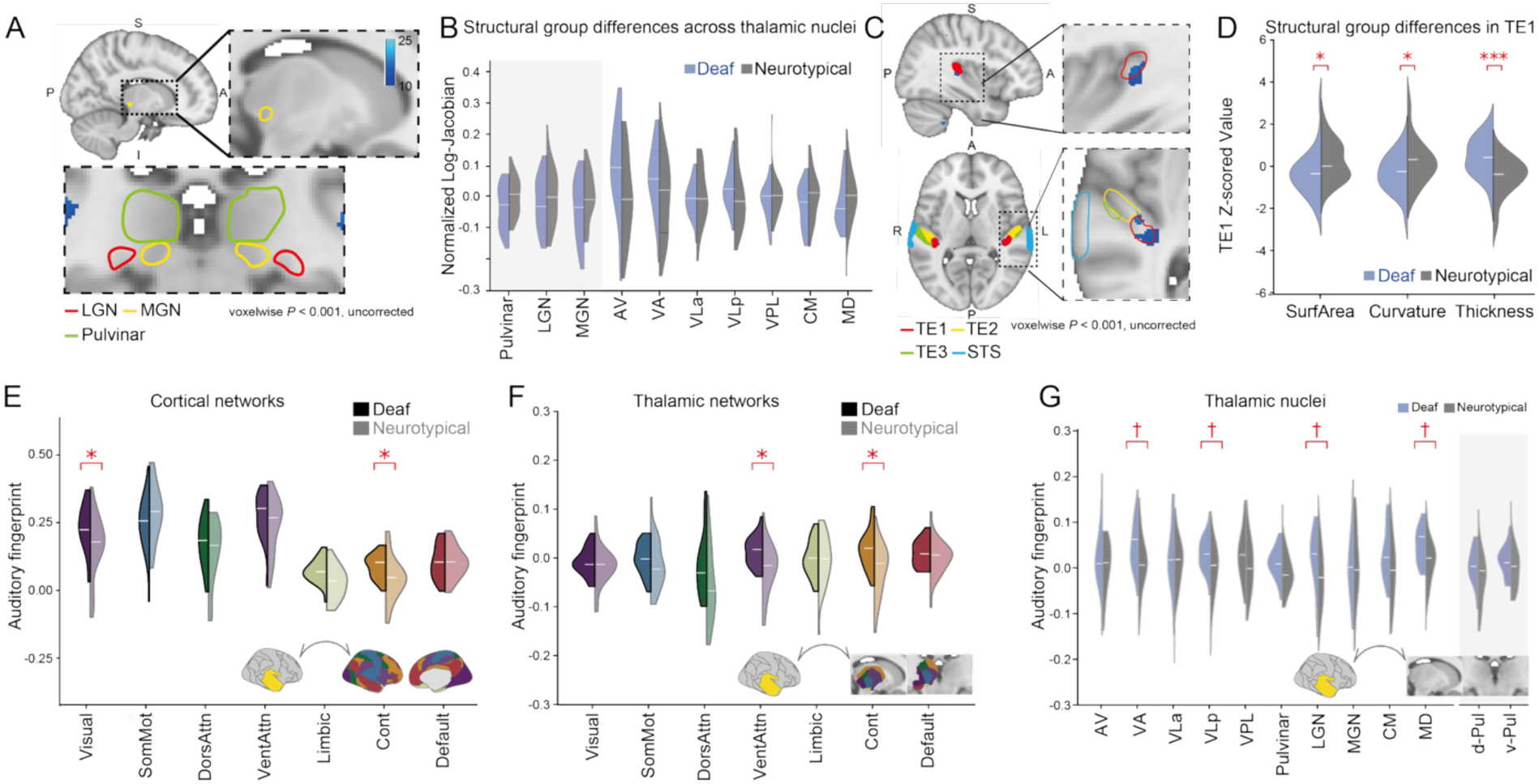
Structural and functional reorganization in early deafness across cortex and thalamus. **(A)** Voxel-wise differences in log-Jacobian values between deaf and neurotypical participants within the thalamus (1,000 permutation tests, *P* < 0.001, uncorrected), highlighted in blue. No significant effects were detected within the medial geniculate nucleus (MGN) or other thalamic nuclei. **(B)** Mean log-Jacobian values within each thalamic nucleus in deaf and neurotypical participants, mean-centered by subtracting the neurotypical group mean for each nucleus. No significant group differences were observed for any thalamic nucleus after FDR correction across 10 nuclei. **(C)** Cortical regions showing significant differences in log-Jacobian between deaf and neurotypical groups (1,000 permutation tests, *P* < 0.001, uncorrected)**. (D)** Structural markers in the primary auditory cortex (TE1) in deaf and neurotypical participants. FDR correction was applied across the three cortical morphometric features. **(E, F)** Functional fingerprint similarity of the cortical auditory network with cortical (E) and thalamic (F) Yeo networks in deaf and neurotypical participants. FDR correction was applied across 7 networks. **(G)** Functional fingerprint similarity between the cortical auditory network and thalamic nuclei. \*\**P* < 0.01 (FDR-corrected), \**P* < 0.05 (FDR-corrected), †*P* < 0.05 (FDR-uncorrected). FDR correction was applied across 10 thalamic nuclei.

We next asked whether the higher-order functional pattern generalized across modality loss. At the cortical level, auditory regions showed increased fingerprint similarity with visual and control network parcels in deaf compared with neurotypical participants (visual: *t*(53) = 2.384, *d* = 0.65, *P* = 0.021, *P*_FDR_ = 0.048; control: *t*(53) = 2.642, *d* = 0.72, *P* = 0.011, *P*_FDR_ = 0.038; Fig. 5E). This pattern remained evident under alternative definitions of auditory cortex (Fig. S12). These results align with prior resting-state and task evidence that early auditory loss reshapes cortical network organization, including recruitment of auditory cortex to visual processing^12,15,67^ and frontoparietal coupling with other large-scale cortical networks^14^. However, the auditory–control effect in deafness was substantially smaller than the visual–control effect in blindness (cf. Fig. 2A; permutation test, *P* = 0.007).

Cross-network plasticity of auditory cortex extended to the thalamus. Auditory cortex showed increased similarity with thalamic ventral attention and control networks (ventral attention: *t*(53) = 2.912, *d* = 0.79, *P* = 0.005, *P*_FDR_ = 0.018; control: *t*(53) = 3.140, *d* = 0.85, *P* = 0.003, *P*_FDR_ = 0.018; Fig. 5F). However, these thalamic effects in deafness were less robust than the thalamic control-related effects observed in blindness. At the level of individual nuclei, auditory cortex showed nominally increased fingerprint similarity with VA, VLp, and MD nuclei (associated with the control network), as well as in the LGN (associated with the visual network) (VA: *t*(53) = 2.421, *d* = 0.66, *P* = 0.019, *P*_FDR_= 0.063; VLp: *t*(53) = 2.612, *d* = 0.71, *P* = 0.012, *P*_FDR_ = 0.058; MD: *t*(53) = 2.675, *d* = 0.72, *P* = 0.010, *P*_FDR_ = 0.058; LGN: *t*(53) = 2.093, *d* = 0.57, *P* = 0.041, *P*_FDR_ = 0.103; Fig. 5G). Thalamic network effects were less robust to a more restrictive auditory ROI definition (visual: *t*(53) = 2.102, *d* = 0.57, *P* = 0.040, *P*_FDR_ = 0.284; Fig. S12C), while parcel-wise analyses localized the group differences primarily to auditory cortical regions (Fig. S13). Complementary analyses anchored on cortical control and visual networks did not reveal significant nucleus-level group differences (Fig. S14A, B).

Together, the blindness and deafness results reveal both common and modality-dependent features of early sensory-loss plasticity. Structural differences in first-order thalamic pathways differed across modality losses, with prominent LGN differences in blindness and no detectable MGN counterpart in deafness. In contrast, visual cortex in blindness and auditory cortex in deafness each showed cross-network functional differences involving distributed cortical networks and control-associated higher-order thalamic systems. The specific network pattern differed by modality, suggesting that higher-order thalamocortical plasticity is a common feature of early sensory loss, while its expression remains shaped by the architecture of the affected sensory system.

## Discussion

Early sensory loss reveals both plasticity constraints and capacities within sensory systems. In blindness, the structural consequences of altered input were concentrated along the primary visual pathway and were organized across visual cortex with respect to V1. Functional differences were distributed more broadly, with visual cortex showing altered relationships with cortical and higher-order thalamic networks that became more pronounced during active nonvisual cognition. Importantly, these structural and functional effects were not spatially independent. Across visual cortex, the strongest cross-network differences occurred where structural effects were greatest, indicating that the architecture of primary sensory input constrains where functional plasticity is expressed. Early deafness showed a related higher-order thalamic pattern despite a different structural expression of sensory loss. Together, these findings suggest that primary sensory pathways constrain how sensory cortex develops, while distributed higher-order thalamocortical systems contribute to the network context in which its functional plasticity is expressed.

### Ascending thalamocortical architecture constrains sensory cortical development

Our structural findings converge with prior evidence that blindness alters the early visual pathway. Human imaging studies have reported reductions in the optic nerves, optic chiasm, optic tracts or radiations, LGN, and visual cortex in congenital or early blindness^39–42,49–53^. Consistent with this work, we found reduced LGN volume and structural differences along the optic tract and optic radiations, together with focal changes in V1 that included reduced surface area and curvature index and increased cortical thickness.

Thalamic structural differences were localized to the LGN, with no detected group differences in the visual pulvinar or other thalamic nuclei. The absence of a macroscopic pulvinar effect is consistent with several studies of blindness or early visual deprivation^51,53,68^. A posterior pulvinar effect has been reported in early blindness, although it occurred within a broad VBM pattern of gray and white matter reductions and became sub-significant when age and sex were included as covariates^39^. Fetal enucleation in macaques can reduce the inferior pulvinar^32^, but this effect may involve subdivisions receiving retinal input that are difficult to resolve with 3T MRI. Altogether, these findings suggest that the dominant thalamic structural effects of blindness are concentrated in nuclei that receive direct retinal input.

The relationship between LGN volume and V1 morphology further suggests that first-order thalamic structure is preferentially associated with some dimensions of cortical anatomy. Within blind participants, LGN volume showed positive associations with V1 surface area and curvature but not cortical thickness. This dissociation fits with developmental accounts in which cortical surface area and folding depend on processes that are partly distinct from those determining cortical thickness^31^. Fetal enucleation studies in macaques provide a developmental precedent for this interpretation, showing that early loss of retinal input reduces LGN size and the extent of striate cortex, alters occipital folding, and produces an atypical cortical territory with mixed striate and extrastriate features (Area X), altering the normal areal and hierarchical relationships within visual cortex^31,32^. Thus, variation in first-order visual thalamic structure is preferentially related to the areal and folding organization of V1, whereas the increased cortical thickness in blindness may reflect other consequences of altered visual experience, including differences in synaptic pruning, dendritic or synaptic architecture, or intracortical myelination^40,41,69–73^.

The magnitude of these structural effects is likely to depend on both the nature and timing of sensory input. Amblyopia provides an informative contrast with early blindness because visual drive is degraded or imbalanced between the eyes, rather than absent bilaterally. Layer-specific changes can occur in the LGN^74,75^, while overall LGN volume remains relatively preserved because input from the non-amblyopic eye is retained. Likewise, early bilateral enucleation in macaques produces larger reductions in LGN and striate cortex than enucleation closer to birth^31,32^. The structural phenotype observed here is therefore best understood as a graded developmental consequence of the amount and timing of ascending sensory input, rather than an invariant feature of blindness.

### Distributed functional plasticity includes higher-order thalamus

The structural consequences of blindness were concentrated near primary sensory input pathways, whereas the functional phenotype involved distributed cortical and thalamic systems. Across cortex, visual regions in blind participants showed greater fingerprint similarity with the control network and reduced similarity with the somatomotor network, consistent with prior studies^7,58,59,76–81^. Within the visual network itself, fingerprint similarity was comparable between groups, consistent with preserved within-network correlation structure and retinotopic-like organization in blindness^59^. Thus, visual cortex retained its within-network functional organization, while group differences were concentrated in its relationships with networks outside the visual system^82^.

These cross-network differences extended to the thalamus, showing that the network context distinguishing visual cortex in blindness spans both cortico-cortical and higher-order thalamocortical systems. Visual cortex showed greater similarity with both control and default-mode thalamic networks and reduced similarity with the thalamic somatomotor network. The default-mode effect may reflect a distinct feature of thalamocortical organization in blindness, although some contribution from blurred boundaries between neighboring control and default-mode territories within the thalamus cannot be excluded. At the level of anatomical nuclei, visual cortex showed increased similarity with several higher-order nuclei, including the AV, VA, VLp, and MD. Cross-network differences were not limited to increased similarity with control-associated thalamic systems. We also found reduced fingerprint similarity between somatomotor cortex and the pulvinar, particularly the ventral pulvinar. Though the higher-order thalamus has received relatively little attention in studies of blindness, a prior study found stronger functional relationships between occipital cortex and ventral lateral and mediodorsal thalamic regions in blindness ^6^. A complementary result from short-term monocular deprivation showed reduced pulvinar connectivity with visual, auditory, and somatomotor cortices^83^. Our findings build on these observations by showing that higher-order thalamic plasticity in blindness is organized at the level of distributed functional networks, paralleling the cross-network differences evident across cortex.

Several pathways could give rise to this correspondence. Many of the nuclei showing stronger similarity with visual cortex are strongly connected with prefrontal cortex and association cortex^84,85^, but lack substantial direct projections to visual cortex. Their functional relationship with visual cortex could therefore arise indirectly through cortical control- or language-related networks. Alternatively, early sensory loss may preserve or reweight developmental connectivity patterns that are attenuated with visual experience. Infant visual cortex shows stronger coupling with prefrontal systems than is observed in sighted adults^86^, raising the possibility that blindness preserves aspects of an early-developing, more exuberantly connected network organization that is typically refined with visual experience. These possibilities are not mutually exclusive, and the fingerprint approach does not distinguish changes originating in cortex from those originating in thalamus.

The spatial organization of the functional effects links distributed plasticity back to the architecture of the sensory system. Visual regions closest to V1 showed the strongest control- and somatomotor-related group differences, and regions with larger reductions in surface area also showed larger cross-network functional differences. This spatial correspondence does not establish that structural differences cause functional plasticity. It does show, however, that the structural and functional phenotypes are organized with respect to a common sensory-system architecture. The same architecture that determines proximity to first-order input therefore also predicts where distributed, cross-network functional differences are most strongly expressed.

Cross-network effects also differed across visual processing pathways, with smaller group differences in dorsal than in ventral and lateral visual cortex. This heterogeneity is consistent with the idea that plasticity operates on pre-existing architecture and latent functional capacities^20^ rather than producing a uniform reassignment of visual cortex. Ventral and lateral regions are typically dominated by visual input but can be recruited during language and tactile processing in blindness^6,87–89^, whereas dorsal regions normally participate in visuomotor and other multisensory functions ^90,91^ ^92^, including in the absence of vision^93^. The weaker dorsal effect may therefore reflect the organization of visual cortex along multiple overlapping axes, including both proximity to the primary cortical entry point of retinal signals and differences in typical network relationships across visual pathways. However, a smaller control-related effect does not imply an absence of dorsal plasticity, as early blindness has been associated with preferential recruitment of right dorsal occipital regions during auditory spatial processing^94^. Developmental differences among these pathways may also contribute. Functional organization in dorsal visual cortex matures earlier than in ventral cortex^95,96^, potentially limiting the extent to which its network organization remains sensitive to postnatal sensory experience. Together, these observations suggest that the capacity for cross-network plasticity is shaped by differences in the typical inputs, network interactions, and developmental timing of visual cortical pathways.

### Higher-order thalamic engagement is enhanced during nonvisual cognition

The functional fingerprint group differences observed at rest became more pronounced during active nonvisual cognition. During the auditory object task, visual-control similarity increased in blind participants relative to rest at both cortical and thalamic levels, whereas visual-somatomotor similarity did not show a comparable task-related increase. The amplification of these higher-order relationships during task engagement suggests that they are functionally relevant to active nonvisual cognition.

Task-dependent expression of cross-modal plasticity has previously been observed primarily at the cortical level^80,97,98^. For example, visual–auditory cross-modal coupling is enhanced during auditory tasks in blind participants relative to rest^98^. Similarly, visual rhythm processing recruits high-level auditory cortex in deaf individuals in a pattern resembling auditory rhythm processing in hearing adults^12^. Our findings extend this state-dependent view to the higher-order thalamus, showing that the same control-related network relationships evident at rest are selectively amplified across both cortical and thalamic levels during active nonvisual processing. Higher-order thalamic involvement therefore appears to be part of the functional network configuration through which visual cortex participates in nonvisual cognition.

#### Shared and modality-dependent features of sensory-loss plasticity

The comparison with early deafness separates features that generalize across early sensory losses from those that depend more strongly on the affected sensory system. The clearest modality difference was structural. Blindness was associated with reduced LGN volume, whereas deafness showed no detectable MGN volume difference (Fig. 6A)^44^. This difference may reflect the organization of the ascending sensory pathways. Retinal signals project directly to the LGN, whereas auditory signals reach the MGN only after multiple brainstem and midbrain relays^99^. The structural consequences of peripheral sensory loss may therefore propagate differently through the two systems. Consistent with this possibility, deaf participants showed evidence of structural differences in the inferior colliculus, which lies closer to the sensory periphery than the MGN. Greater heterogeneity in deafness, including variability in etiology and residual auditory input, may also reduce sensitivity to MGN effects.

**Figure 6.**
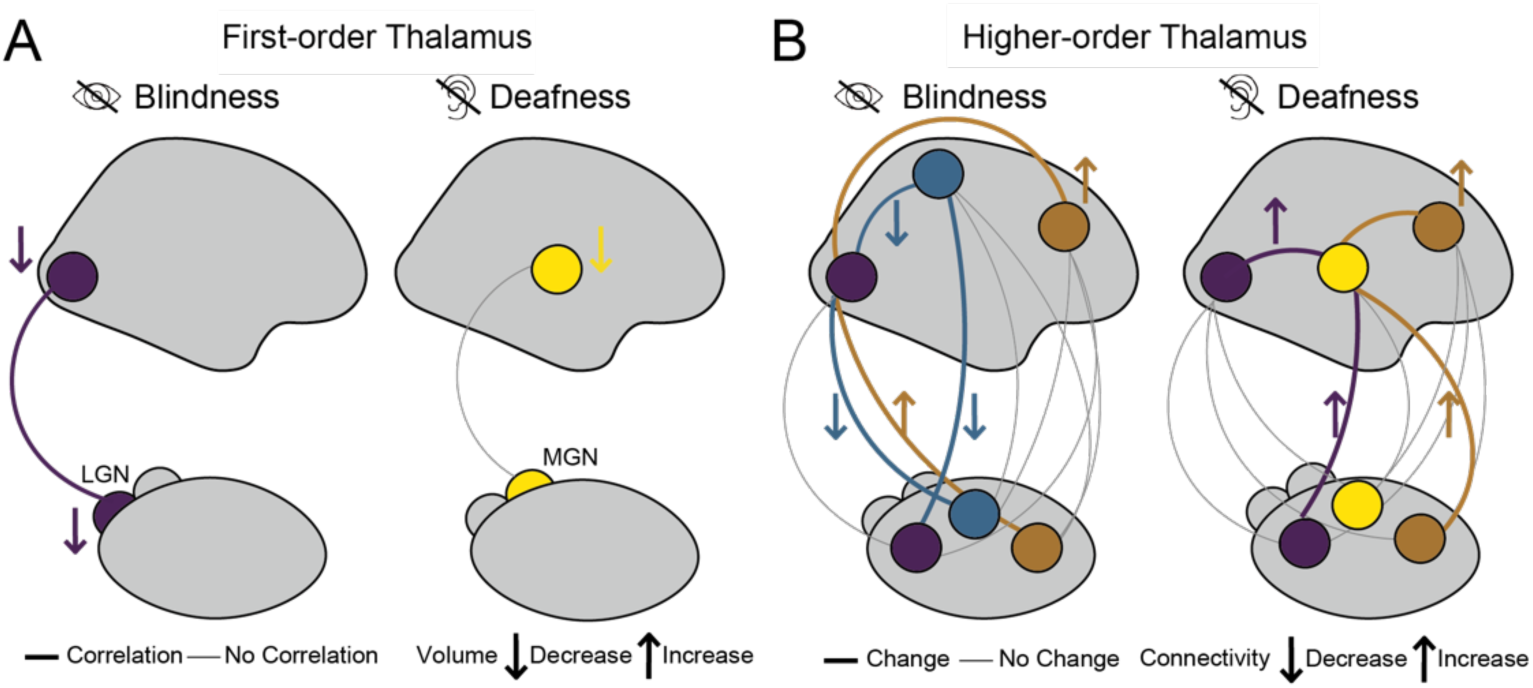
Summary of thalamic involvement in cross-modal reorganization. **(A)** In blindness, LGN volume and V1 surface area are both reduced and correlated across individuals. In contrast, deafness is not associated with detectable MGN volume loss, and A1 surface area reduction is weaker than V1 reduction in blindness. **(B)** In blindness, the visual cortex shows increased fingerprint similarity with the control network and decreased similarity with the somatomotor network at both the cortical and higher-order thalamic levels. In deafness, the auditory cortex shows increased similarity with the visual and control networks at both levels.

Cortical structure showed both common and modality-dependent features. Primary visual and auditory cortex showed changes in the same overall direction, including increased cortical thickness, while the reduction in surface area was larger in blindness. Surface area may therefore be more tightly coupled to the architecture and developmental influence of ascending sensory pathways, whereas increased cortical thickness may reflect consequences of atypical sensory experience that operate more broadly across sensory cortices, potentially arising from experience-dependent differences in pruning, intracortical maturation, dendritic or synaptic architecture, or myelination^40,41,69–71,100^.

Despite these differences in first-order structural phenotype, blindness and deafness showed a common pattern of cross-network functional plasticity across distributed cortical and higher-order thalamic networks (Fig. 6B). In both sensory-loss groups, sensory cortex associated with the absent modality showed increased functional alignment with networks outside its canonical sensory system, including control-related networks consistent with prior work^6,81,101–103^. This commonality extended to higher-order thalamus, with control-related differences evident at both cortical and thalamic levels. Although the precise network pattern differed by modality and nucleus-level effects were less robust in deafness, the convergence of control-related cortical and thalamic effects suggests that higher-order thalamocortical plasticity is a common feature of early sensory loss, whose specific expression remains shaped by the architecture of the affected sensory system.

#### Limitations

Several limitations should be considered when interpreting these findings. First, functional fingerprint analyses can identify sensory-loss-related differences in thalamocortical functional organization but cannot determine the circuit-level mechanisms that give rise to those differences. Because these functional profiles reflect functional correspondence between cortex and thalamus, changes at either level could contribute to the observed thalamocortical differences. Thus, the present findings identify coordinated functional differences across cortical and thalamic systems, but they do not determine whether those differences arise from changes in cortical organization, thalamic organization, the weighting of pre-existing pathways, or reciprocal interactions between these systems^17,20^. Dissecting these mechanisms will require animal models and causal circuit manipulations that can directly test how changes shape functional network organization after early sensory loss. Second, the functional MRI data were acquired at 3-mm isotropic resolution, limiting the precision within small thalamic nuclei. Although effects were evident at both network and nucleus levels, partial-volume effects, imperfect nucleus localization, and measurement noise may influence nucleus-level estimates. Higher-resolution imaging with individualized thalamic segmentation may help resolve finer thalamic details of these cross-network differences. Finally, the study included modest sample sizes and typical variability in the onset, duration, and etiology of sensory loss. This heterogeneity likely limits our sensitivity to determine how specific developmental histories influence thalamocortical organization. Larger cohorts with more detailed clinical characterization will be needed to distinguish the effects of onset, residual sensory experience, etiology, and duration.

Nevertheless, many of the structural and functional patterns observed here converge with prior findings in independent samples, suggesting that the main effects reflect reliable features of early and early sensory loss.

### Conclusion

Thalamocortical architecture may therefore shape both constraints and capacities for plasticity after early sensory loss. Structural consequences remain closely tied to the organization of ascending sensory pathways, while functional reorganization extends across distributed cortical and higher-order thalamic networks. The common higher-order involvement observed in blindness and deafness, together with modality-dependent differences in structural and network expression, suggests that plasticity emerges from an interaction between altered sensory experience and the pre-existing architecture of each sensory system. These findings may help explain why some features of cortical organization remain closely tied to early sensory pathways, while others retain substantial capacity for functional reorganization.

## STAR★METHODS

### KEY RESOURCES TABLE

|  |  |  |
| --- | --- | --- |
| **Biological samples** |  |  |
| Early blind and neurotypical participant MRI data | Xu et al., 2023<sup>66</sup> | https://doi.org/10.1371/journal.pbio.3001930. |
| Early deaf and neurotypical participant MRI data | Bonna et al., 2021<sup>14</sup> | https://doi.org/10.1007/s11682-020-00346-y. |
| **Deposited data** |  |  |
| Analyzed data | This paper | https://zenodo.org/records/21456505. |
| Code | This paper | https://github.com/monami-nishio/blind_deaf_thalamus |
| **Software and algorithms** |  |  |
| fMRIPrep v25.1.4 | Esteban et al., 2019<sup>104</sup> | RRID:SCR_016216<br>https://fmriprep.org/en/stable/ |
| FreeSurfer v7.3.2 | Fischl et al., 2012<sup>105</sup> | RRID:SCR_001847<br>https://surfer.nmr.mgh.harvard.edu/ |
| ANTs v2.6.2 | Avants et al., 2011<sup>106</sup> | RRID:SCR_004757<br>https://github.com/antsx/ants |
| FSL | Jenkinson et al., 2012<sup>107</sup> | RRID:SCR_002823<br>https://fsl.fmrib.ox.ac.uk/fsl/docs/ |
| AFNI | Cox, 1996<sup>108</sup> | RRID:SCR_005927<br>https://afni.nimh.nih.gov/ |
| THOMAS | Su et al., 2019<sup>48</sup> | https://doi.org/10.1016/j.neuroimage.2019.03.021 |
| tvb-gdist | Mitchell et al., 1987<sup>109</sup> | https://github.com/the-virtual-brain/tvb-gdist |
| **Atlases and maps** |  |  |
| Schaefer 400-parcel cortical atlas | Schaefer et al., 2018<sup>65</sup> | https://doi.org/10.1093/cercor/bhx179<br>https://github.com/ThomasYeoLab/CBIG/tree/master/stable_projects/brain_parcellation/Schaefer2018_LocalGlobal |
| Yeo 7-Network cortical parcellation | Yeo et al., 2011<sup>57</sup> | https://doi.org/10.1152/jn.00338.2011 |
| Yeo thalamic functional parcellation | Yeo et al., 2011<sup>57</sup> / associated resources | https://www.dropbox.com/scl/fi/sgo2yqds3kmgja1opr0lu/ThalamusParcellation.zip?rlkey=vyqlldolsq42q7acro6mfsodd&e=1&dl=0 |
| Wang probabilistic retinotopic atlas | Wang et al., 2015<sup>55</sup> | https://doi.org/10.1093/cercor/bhu277 |
| Julich-Brain cytoarchitectonic atlas | Amunts et al. 2020<sup>110</sup> | https://doi.org/10.1126/science.abb4588 |
| Sensory-Association (S-A) axis | Sydnor et al., 2021<sup>111</sup> | https://doi.org/10.1016/j.neuron.2021.06.016<br>https://github.com/PennLINC/S-A_ArchetypalAxis |

### EXPERIMENTAL MODEL AND SUBJECT DETAILS

#### Participants

##### Early Blind

Data from early blind and matched neurotypical participants were obtained from Xu et al., 2023^66^. The dataset initially included 16 early blind participants and 20 neurotypical participants. One early blind participant was excluded because the required anatomical imaging data were unavailable, and one neurotypical participant was excluded due to resting-state fMRI preprocessing failure. The final sample comprised 15 early blind participants (*M* age = 32.8 years, *SD* = 4.5) and 19 neurotypical participants (*M* age = 32.5 years, *SD* = 5.9). The original study reported that the groups did not differ in age and sex. The etiology of blindness was mainly ocular or retinal, with 2 cases including optic nerve. The early blind group reported, at most, faint light perception and had no visual memories. All blind participants were blind from birth, except three participants who had visual difficulties from birth and fully lost vision at 8 months, 2 years, and 4 years. This study was conducted according to the principles expressed in the Declaration of Helsinki. The ethical committee of the University of Trento approved the experimental protocol in this study (protocol 2014–007).

##### Early Deaf

Data from early deaf and neurotypical participants were obtained from Bonna et al., 2021^14^. The dataset initially included 28 early deaf participants and 30 neurotypical participants. Three early deaf participants were excluded because of image acquisition errors, resulting in 25 early deaf participants (13 females; age: *M* = 29.4; *SD* = 6.04) and 30 neurotypical participants (18 females; age: *M* = 26.4; *SD* = 4.51). The groups did not differ in age and sex. The etiology of deafness was either genetic (hereditary deafness) or pregnancy-related (maternal disease or drug side effects). The mean hearing loss was 100.2 dB (range 70–120 dB) for the left ear and 101.4 dB (60–120 dB) for the right ear. All subjects had some experience with hearing aids (currently or in the past), but did not rely on them on a daily basis. Based on a self-assessment survey, all subjects were proficient users of Polish Sign Language. The ethical committee of Jagiellonian University approved the experimental protocol in this study.

### METHOD DETAILS

#### MRI acquisition

##### Early Blind

MRI data were acquired using a 3T MAGNETOM Prisma scanner (Siemens, Erlangen, Germany) equipped with a 64-channel head–neck coil at the Center for Mind/Brain Sciences, University of Trento. Resting-state functional images were acquired using a simultaneous multislice echo-planar imaging (EPI) sequence. The acquisition plane was parallel to the bicommissural plane, with phase encoding from anterior to posterior (TR = 1000 ms; TE = 28 ms; FA = 59°; multiband factor = 5; FOV = 198 mm; matrix size = 66 × 66; 65 axial slices; slice thickness = 3 mm; voxel size = 3 × 3 × 3.3 mm). Each resting-state run lasted 8 minutes. Resting-state scanning was followed by 10 task runs, each lasting 5 min 30 s, during which participants performed word verification tasks based on auditorily presented stimuli. T1-weighted anatomical images were acquired using a MPRAGE sequence (TR = 2140 ms; TE = 2.9 ms; FA = 12°; FOV = 288 mm; 208 sagittal slices; voxel size = 1 × 1 × 1 mm).

##### Early Deaf

MRI data were acquired using a 3T MAGNETOM Tim Trio scanner (Siemens, Erlangen, Germany) equipped with a 32-channel head coil. Resting-state functional images were acquired using a gradient-echo EPI sequence (TR = 2190 ms; TE = 30 ms; FA = 90°; FOV = 192 mm; matrix size = 64 × 64; 33 axial slices; slice thickness = 3.6 mm; voxel size = 3 × 3 × 3.6 mm). Each resting-state run lasted 10 minutes. T1-weighted anatomical images were acquired using a MPRAGE sequence (TR = 2530 ms; TE = 3.32 ms; FA = 7°; FOV = 256 mm; 176 sagittal slices; voxel size = 1 × 1 × 1 mm).

#### MRI preprocessing

T1-weighted anatomical images and functional MRI time series from the blind and deaf datasets were preprocessed using fMRIPrep 25.1.4^104^.

For anatomical preprocessing, T1-weighted images were corrected for intensity non-uniformity using N4BiasFieldCorrection from ANTs 2.6.2, skull-stripped using the ANTs brain extraction workflow implemented in Nipype 1.10.0, and segmented into brain tissue classes using FSL FAST. Brain surfaces were then reconstructed using FreeSurfer 7.3.2. Volume-based spatial normalization of the T1-weighted image, one standard space (MNI152NLin2009cAsym), was performed through nonlinear registration with ANTs.

For functional preprocessing, a skull-stripped reference BOLD image was generated for each run. A B0 field map was estimated from two or more echo-planar imaging (EPI) reference runs, with top-up from FSL, and aligned to the target EPI reference run via rigid registration. The BOLD reference was co-registered with rigid transformations (six degrees of freedom) to the T1-weighted reference using bbregister in FreeSurfer. Head-motion parameters were estimated with respect to the BOLD reference before spatiotemporal filtering using FSL’s mcflirt. BOLD runs were slice-time corrected using 3dTshift from AFNI 20160207 and resampled to their original native space by applying a single composite transform to correct for head motion and susceptibility distortions. The estimated head-motion parameters were subsequently regressed from the BOLD time series as nuisance covariates. The BOLD time series were also resampled onto the fsaverage surface and into standard space, generating a preprocessed BOLD run in MNI152NLin2009cAsym space.

#### Task paradigm

Details of the task paradigm are reported in Xu et al., 2023^66^. Briefly, the task was designed to dissociate shape-based and conceptual representations of object words in sighted and early blind participants. Stimuli consisted of Italian words referring to everyday manmade objects. The stimulus set was designed to dissociate shape-based and conceptual representations while restricting items to a single taxonomic category. This design allowed responses to object-word stimuli to be compared between sighted and early blind participants while minimizing differences attributable to broad category structure. The stimulus set included 21 object words. Auditory versions of the words were recorded by a professional narrator, trimmed for silence, and intensity-normalized to 70 dB using the open-source program Praat. In the present study, these task data were used for imaging-based analyses, and behavioral ratings of object properties were not analyzed.

#### Cortical and Thalamic Parcellation

Cortical analyses were conducted using the Schaefer 400-parcel atlas^65^, which is based on Yeo 7 functional networks^57^. Visual and auditory cortical parcels were defined based on overlap with the Wang atlas^55^ and the Julich atlas^110^, respectively. This procedure identified 86 visual parcels across the two hemispheres, which were retained as separate parcels for parcel-wise structural and functional analyses. Schaefer parcels exhibiting at least 5% overlap with either atlas were classified as visual or auditory parcels, respectively. Schaefer parcels overlapping Wang atlas visual regions were further grouped into four processing streams according to their Wang labels: primary (V1–V3), ventral (hV4– PHC2), lateral (LO1–2 and TO1–2), and dorsal (V3A, V3B, and IPS0–5). This procedure enabled analyses of visual and auditory cortices within a common cortical parcellation framework. For network-level analyses, Schaefer parcels were assigned to the Yeo 7-network parcellation. To subdivide the somatomotor network into auditory and non-auditory regions, parcels assigned to the somatomotor network that overlapped with the Julich atlas were classified as auditory, whereas those with no overlap were classified as non-auditory.

Thalamic analyses used both anatomical and functional parcellations. Anatomically defined thalamic nuclei were identified using the THOMAS atlas, a multi-atlas label fusion method for segmenting individual thalamic nuclei from anatomical MRI data^48^. Functionally defined thalamic territories were identified using the Yeo thalamic parcellation, which was derived from voxel-wise functional connectivity patterns between the thalamus and cortex. These thalamic parcellations were used to compare effects across individual thalamic nuclei and across thalamic territories associated with large-scale functional networks.

#### Jacobian Deformation

To assess local volumetric differences between groups, we performed a Jacobian deformation analysis^47^. Individual T1-weighted anatomical images were nonlinearly registered to MNI template, and the log-transformed Jacobian determinant of the deformation field was calculated at each voxel, providing a measure of local expansion or contraction relative to the template. Group differences were assessed separately for the blind–neurotypical and deaf–neurotypical comparisons using nonparametric permutation tests. Group labels were randomly shuffled 1,000 times, and the voxel-wise group difference was recalculated for each permutation. Statistical significance was determined at the voxel level with a threshold of *P* < 0.001, uncorrected. For nucleus-level analyses, mean log-transformed Jacobian deformation values were extracted within each THOMAS-defined thalamic nucleus and compared between groups using two-tailed independent-samples t-tests, with FDR correction across the 10 nuclei. For visualization in Figures 1B and 5B, participant-level values were mean-centered within each nucleus by subtracting the corresponding neurotypical group mean.

Voxel-wise group differences in log-Jacobian maps were assessed using nonparametric permutation testing. The observed difference in mean log-Jacobian values between groups was compared with a null distribution generated by randomly permuting group labels 1,000 times while preserving group sizes. Two-tailed voxel-wise P values were calculated as the proportion of permutations yielding an absolute group difference at least as large as the observed absolute difference, with voxels considered significant at P < 0.001 (uncorrected).

#### Thalamic nucleus volume estimation

Thalamic nuclei were segmented from each participant’s T1-weighted anatomical image using THOMAS, an automated multi-atlas-based method for probabilistic thalamic parcellation^48^. For each participant, the volume of each THOMAS-defined nucleus was extracted and normalized to the mean volume of that nucleus in the corresponding neurotypical group, providing a relative measure of nucleus volume with respect to the neurotypical reference mean. Group differences in normalized nucleus volumes were then assessed by comparing sensory-loss (early blind or early deaf) and neurotypical participants. For the pulvinar region-specific analyses, pulvinar ROIs defined in MNI152NLin2009cAsym standard space were transformed into each participant’s native T1-weighted space using the participant-specific nonlinear transformation estimated by fMRIPrep. The volume of each pulvinar parcel was then calculated from the number of non-zero voxels within the transformed ROI and the corresponding native-space voxel dimensions.

#### Cortical structural feature extraction

Cortical structural features were extracted from high-resolution T1-weighted MRI scans using FreeSurfer 7.3.2 (http://surfer.nmr.mgh.harvard.edu). Following surface reconstruction, cortical morphometric features were extracted on each participant’s native cortical surface. Surface area, cortical thickness, and curvature index were computed and summarized within atlas-defined regions. Visual regions were defined using the Wang atlas^55^, auditory regions using the Julich atlas^110^, and whole-brain analyses were performed using the Schaefer 400-parcel atlas^65^.

#### Functional fingerprint analysis

Functional fingerprint analyses^54^ were used to quantify the similarity between distributed cortical and thalamic networks. For network-level analyses in blindness, sensory cortex was defined as the Yeo visual network. For parcel-wise spatial analyses across visual cortex, including the analyses of geodesic distance and structure–function correspondence in Figures 3 and S9, functional differences were quantified separately across the 86 Schaefer parcels overlapping Wang atlas visual regions. For analyses in deafness, sensory cortex was defined as an auditory cortical ROI comprising Schaefer parcels assigned to the Yeo somatomotor network that overlapped the Julich auditory atlas. We used this approach to test whether sensory cortex showed altered functional coupling with large-scale cortical networks and with thalamic territories associated with those networks.

For each participant, mean resting-state time series were extracted from each of the 400 Schaefer cortical parcels. Pairwise Spearman correlations were computed between all cortical parcels to generate a parcel-wise functional coupling matrix. Each cortical parcel was then characterized by its functional fingerprint, defined as its profile of functional coupling with all other cortical parcels. We then computed a temporal correlation matrix of thalamic voxels by cortical parcel. For each thalamic voxel, the voxel time series was correlated with the mean time series of each Schaefer cortical parcel. Each row of this matrix represented the cortical correlation profile of one thalamic voxel. To quantify fingerprint similarity, each thalamic voxel correlation profile was spatially correlated with each Schaefer parcel correlation profile. This produced a thalamic voxel-by-cortical parcel spatial correlation fingerprint matrix.

To assess relationships with large-scale functional networks, thalamic voxels were grouped according to the Yeo thalamic parcellation, and cortical parcels were grouped according to their Yeo 7 network assignments. Spatial correlation fingerprint values were averaged across thalamic voxels and cortical parcels within each network pair to estimate thalamic-to-cortical network fingerprint similarity for each participant. To assess relationships with individual thalamic nuclei, thalamic voxels were grouped according to THOMAS-defined nuclei, and spatial correlation fingerprint values were averaged between each thalamic nucleus and the cortical parcels of interest.

For analyses of task-state functional fingerprints in the blindness dataset, the same procedure was applied to the preprocessed task BOLD time series. Task-evoked responses were not regressed from the time series before calculation of the functional coupling matrices; thus, task-state fingerprints reflected the overall correlation structure during task performance, including task-evoked components.^65^

#### Geodesic distance from sensory regions

Geodesic distance along the cortical surface was calculated using tvb-gdist^109^ module that approximates the shortest path between two nodes on a triangular surface mesh. We selected parcels in the Schaefer 400 parcellation^65^ corresponding to the V1 of Wang atlas and TE1 of Julich atlas. To reduce computational demands while maintaining spatial coverage, vertices within each region were partitioned using k-means clustering, with the number of clusters (k) set to 5% of the total number of vertices in the region (rounded down to the nearest integer). One vertex was randomly selected from each cluster as a seed node, resulting in approximately 5% of the regional vertices being used as spatially distributed seeds. Each cortical node was then assigned a distance value based on the minimum geodesic distance along the “midthickness” surface to any of the seed nodes. Finally, the average of the minimum geodesic distances for nodes within each cortical parcel of the Schaefer 400 parcellation was calculated.

#### Sensorimotor–Association (S-A) axis

We used the S-A axis derived by Sydnor and colleagues^111^. This map integrates various cortical hierarchies, including functional connectivity gradients, evolutionary cortical expansion patterns, anatomical ratios, allometric scaling, brain metabolism measures, perfusion indices, gene expression patterns, primary modes of brain function, cytoarchitectural similarity gradients, and cortical thickness.

### QUANTIFICATION AND STATISTICAL ANALYSIS

Statistical analyses were performed in Python unless otherwise specified. Group differences in continuous measures were assessed using two-tailed independent-samples t-tests and are reported with Cohen’s d alongside t and P values. Pearson correlation coefficients were used to assess relationships among structural and functional measures within each group and are reported with 95% confidence intervals. Where correlations were compared between groups, differences were tested directly using Fisher r-to-z tests. For comparisons of functional fingerprint measures between resting-state and task conditions, group × condition interactions were assessed using mixed-design analyses of variance, with condition as a within-subject factor and group as a between-subject factor. Within-group rest–task differences were subsequently assessed using two-tailed paired-samples t-tests and are reported with Cohen’s d*_z_*. Group × stream interactions across visual processing streams were assessed using mixed-design analyses of variance, with stream as a within-subject factor and group as a between-subject factor.

Direct comparisons of sensory-loss effects between the blindness and deafness datasets were assessed using nonparametric permutation tests. For each permutation, group labels were randomly shuffled within each dataset while preserving group sizes. The sensory-loss versus neurotypical group difference was then recalculated separately for each dataset, and the difference between these two group effects was computed. This procedure was repeated 1,000 times to generate a null distribution of between-dataset differences. Two-tailed P values were calculated as the proportion of permutations yielding an absolute between-dataset difference at least as large as the observed absolute difference.

To control for multiple comparisons, P values were adjusted using the Benjamini–Hochberg false discovery rate (FDR) procedure. FDR correction was applied separately within each family of related statistical tests, including (i) comparisons across the 10 thalamic nuclei, (ii) comparisons across the three cortical morphometric features (surface area, cortical thickness, and curvature), (iii) comparisons across 7 or 8 networks, and (iv) multiple pairwise comparisons, as appropriate. For the direct comparison of cortical morphometric effects between blindness and deafness, permutation-test P values were FDR-corrected across surface area, curvature, and cortical thickness. Voxel-wise Jacobian analyses were used to provide a spatially resolved assessment of structural differences without constraining effects to predefined anatomical regions and were thresholded at P < 0.001, uncorrected for multiple comparisons.

Relationships among cortical structural and functional difference maps, geodesic distance from primary sensory cortex (V1 or TE1), and S–A axis position were quantified using Pearson correlation coefficients. Correlations involving cortical maps were additionally evaluated using spatial permutation (spin) tests^112^ where appropriate to account for spatial autocorrelation. Unless otherwise specified, statistical significance was defined as P < 0.05; for analyses subjected to FDR correction, significance was determined using the FDR-adjusted P value. Reported degrees of freedom were calculated as n₁ + n₂ − 2 for independent-samples t-tests, n − 1 for paired-samples t-tests, and n − 2 for Pearson correlations.

## Acknowledgements

This work was supported by an NIH grant P50MH132642 (to M.J.A.). M.N. was supported by Quad Fellowship and Nakajima Foundation Scholarship. The funders had no role in the study design, data collection and interpretation, or the decision to submit the work for publication.

## Author contributions

Conceptualization, M.N., X.L., and M.J.A.; Methodology, M.N., X.L., and M.J.A.; Formal analysis, M.N.; Resources and data acquisition, Y.X., M.Z., M.S., and O.C.; Writing – original draft, M.N. and M.J.A.; Writing – review & editing, all authors; Supervision, A.P.M. and M.J.A.; Funding acquisition, M.J.A.

## Supplementary Figures

**Supplementary Figure 1.**
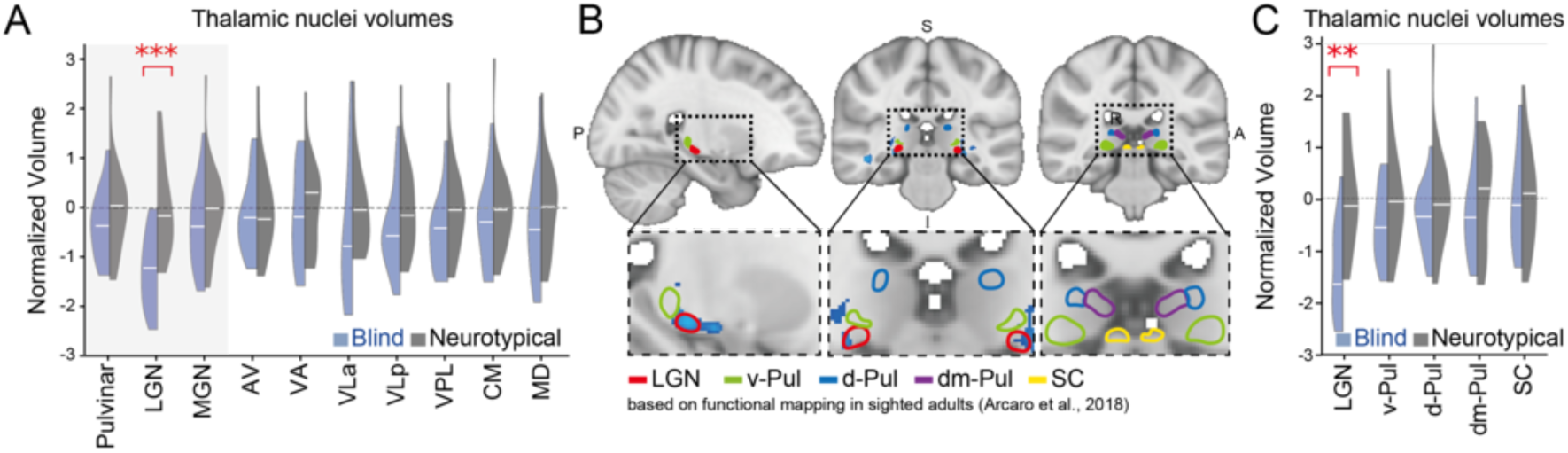
Thalamic structural reorganization in early sensory loss. **(A)** Anatomically-defined thalamic nucleus volumes in blind participants and neurotypical controls. For visualization across nuclei with substantially different absolute volumes, volumes were z-scored separately within each nucleus. Visual- and auditory-related nuclei are highlighted in gray. FDR correction was applied across 10 thalamic nuclei. **(B)** Voxels showing significantly higher Jacobian values in blind compared with neurotypical participants in relation to the LGN, subdivisions of the pulvinar, and SC based on prior functional mapping in sighted adults^54^. **(C)** Functionally-defined volumes for LGN, subdivisions of the pulvinar, and SC in blind and control participants. Significance: \*\*\**P* < 0.001, \*\**P* < 0.01, \**P* < 0.05. FDR-corrected. FDR correction was applied across the five thalamic subdivisions.

**Supplementary Figure 2.**
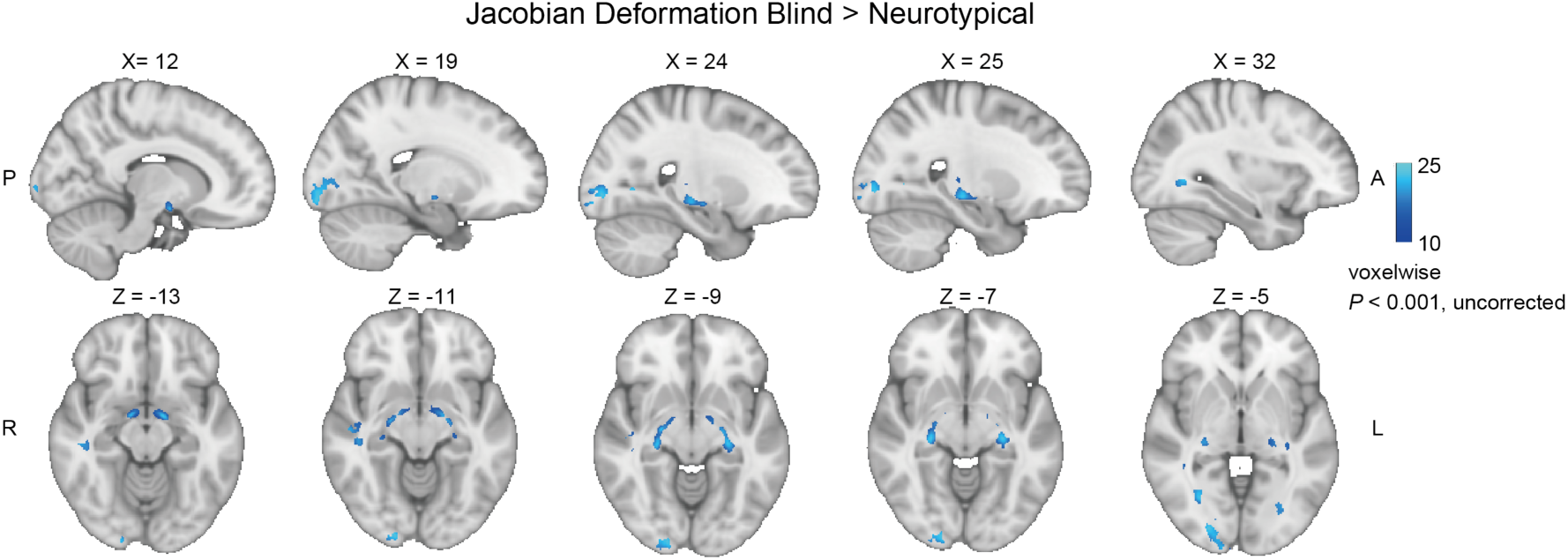
Jacobian deformation of white matter tract in early blindness. Voxels showing significantly higher log-Jacobian values in blind compared with neurotypical participants are shown in blue (1,000 permutation tests, P < 0.001). Log-Jacobian values were derived from deformation to the MNI template, with higher values indicating greater local expansion during registration and therefore smaller native anatomy relative to the template. Significant effects were evident along the optic tract and optic radiations.

**Supplementary Figure 3.**
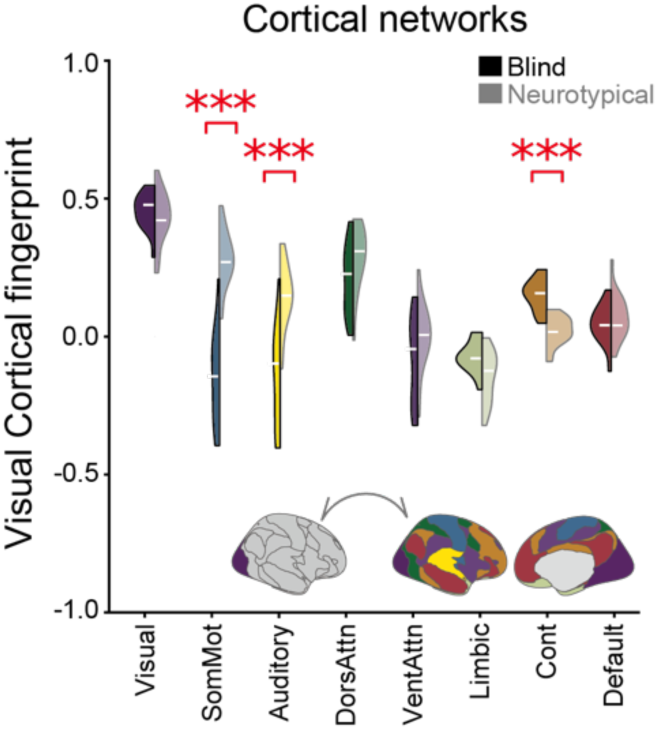
Cortical functional reorganization in early blindness. Functional fingerprint similarity of cortical visual network to cortical Yeo networks in blind and neurotypical groups. \*\*\**P* < 0.001 (FDR-corrected), \*\**P* < 0.01 (FDR-corrected), \**P* < 0.05 (FDR-corrected). FDR correction was applied across eight networks.

**Supplementary Figure 4.**
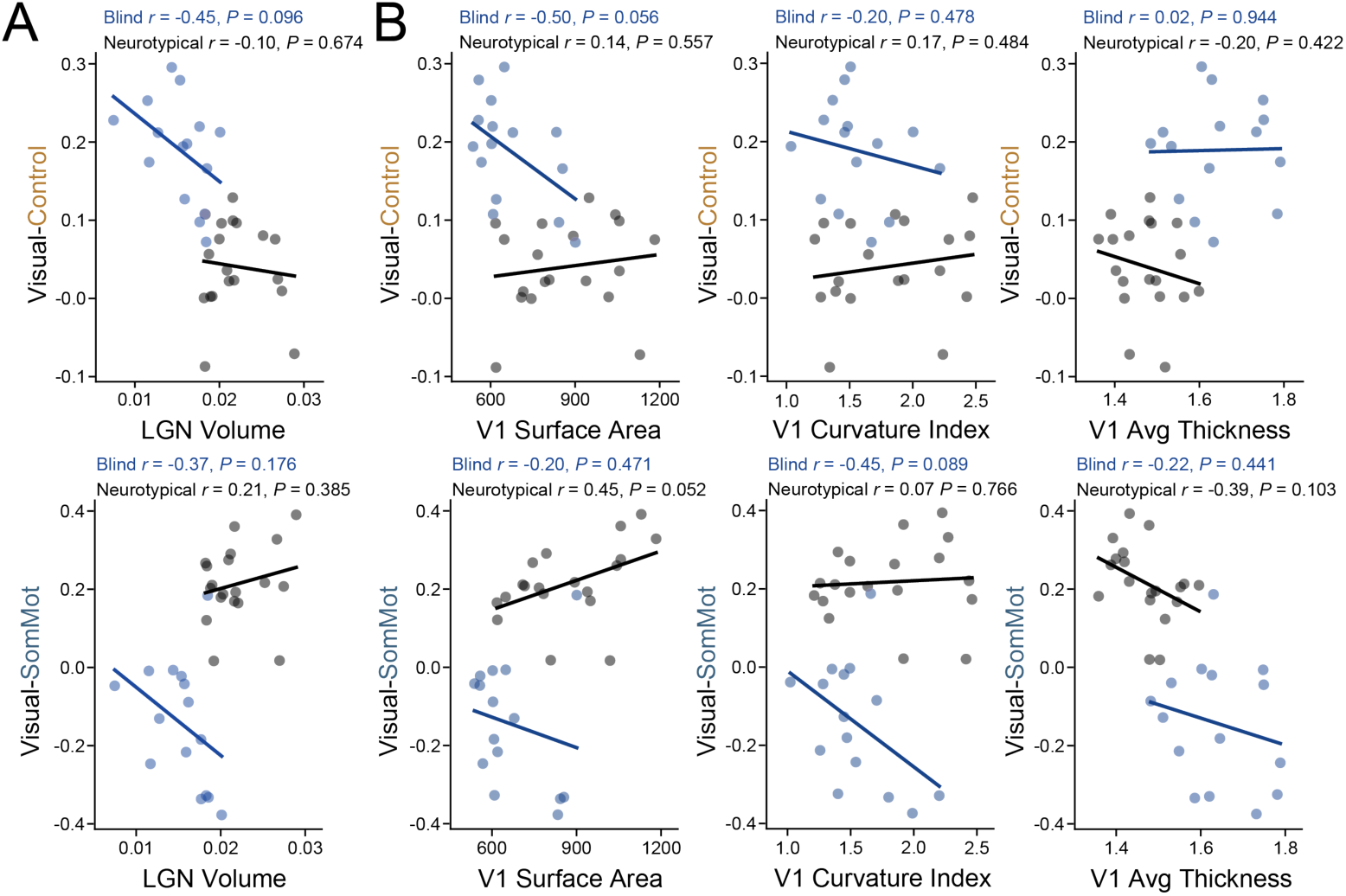
Correlation between structural and functional reorganization. Correlations of **(A)** LGN volume and **(B)** V1 structural features with functional fingerprint similarity. Top: visual–control fingerprint similarity. Bottom: visual–somatomotor fingerprint similarity.

**Supplementary Figure 5.**
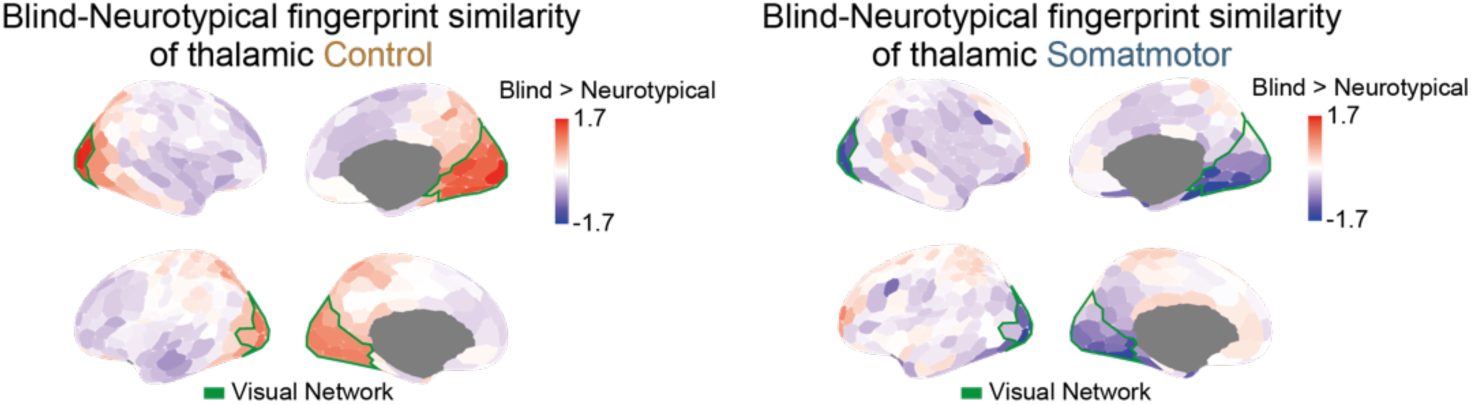
Thalamocortical functional reorganization in early blindness. Group differences between early blind and neurotypical participants in the fingerprint similarity between the thalamic control and somatomotor networks and cortical parcels. Cortical parcels overlapping with the visual network are highlighted in green.

**Supplementary Figure 6.**
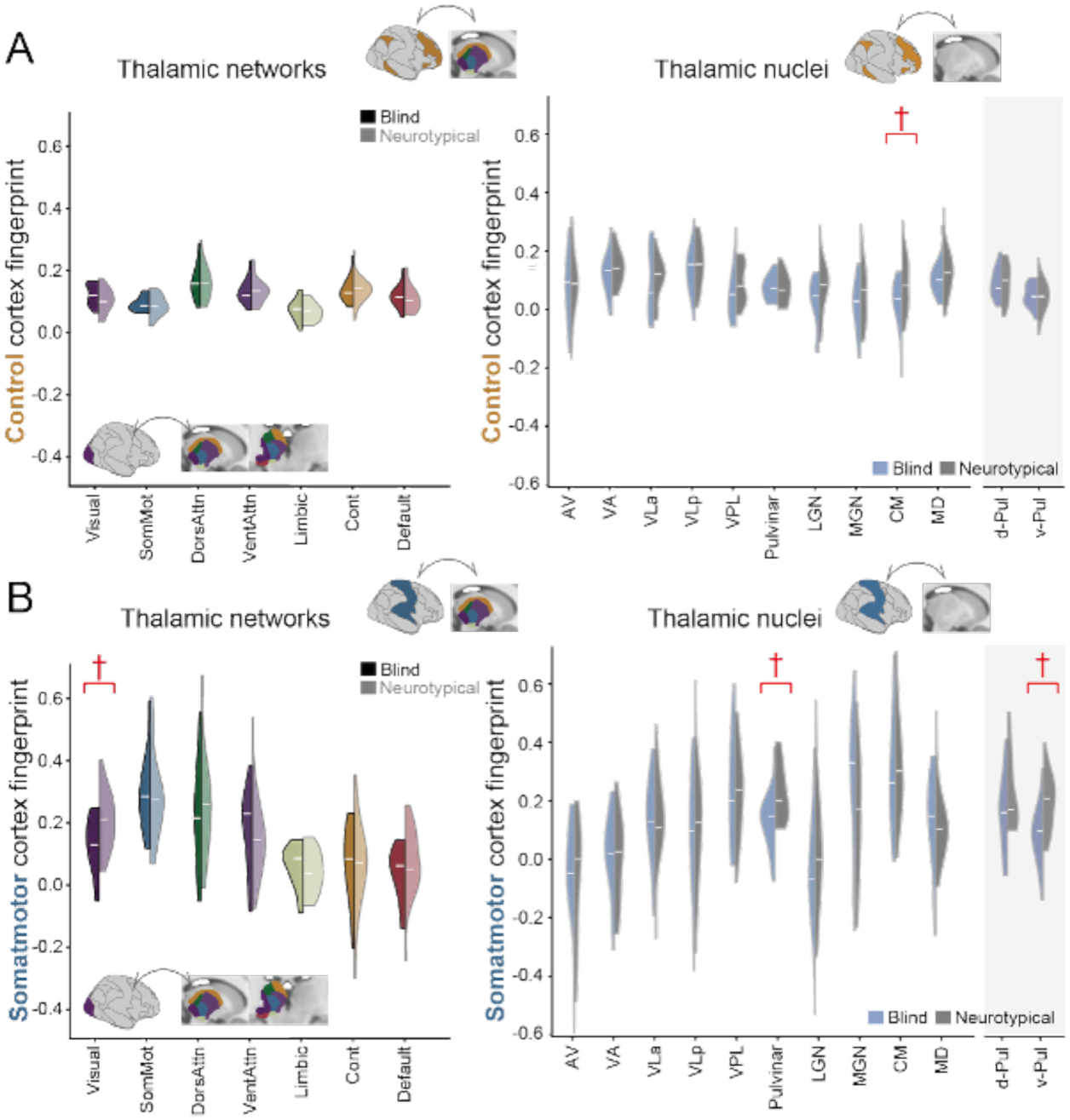
Thalamic functional fingerprint plasticity in early blindness. **(A, B)** Functional fingerprint similarity of the cortical (A) control and (B) somatomotor networks and thalamic nuclei in blindness. (A) The cortical control network showed a difference in fingerprint similarity with the centromedial (CM) nucleus, although this effect did not survive FDR correction. (B) Mirroring the decreased fingerprint similarity between the cortical visual network and thalamic somatomotor regions, the cortical somatomotor network showed reduced fingerprint similarity with the thalamic visual network and the pulvinar (visual network *t*(32) = −2.375, *P* = 0.023, *P*FDR = 0.165, Pul *t*(32) = −2.764, *P* = 0.009, *P*FDR = 0.094). An exploratory analysis of pulvinar subdivisions suggested that this effect was strongest in the ventral pulvinar (vPul *t*(32) = −3.570, *P* = 0.002; dPul *t*(32) = −1.516, P = 0.139). †*P* < 0.05 (FDR-uncorrected). FDR correction was applied separately across the 7 cortical networks for network-level analyses and across the 10 thalamic nuclei for nucleus-level analyses.

**Supplementary Figure 7.**
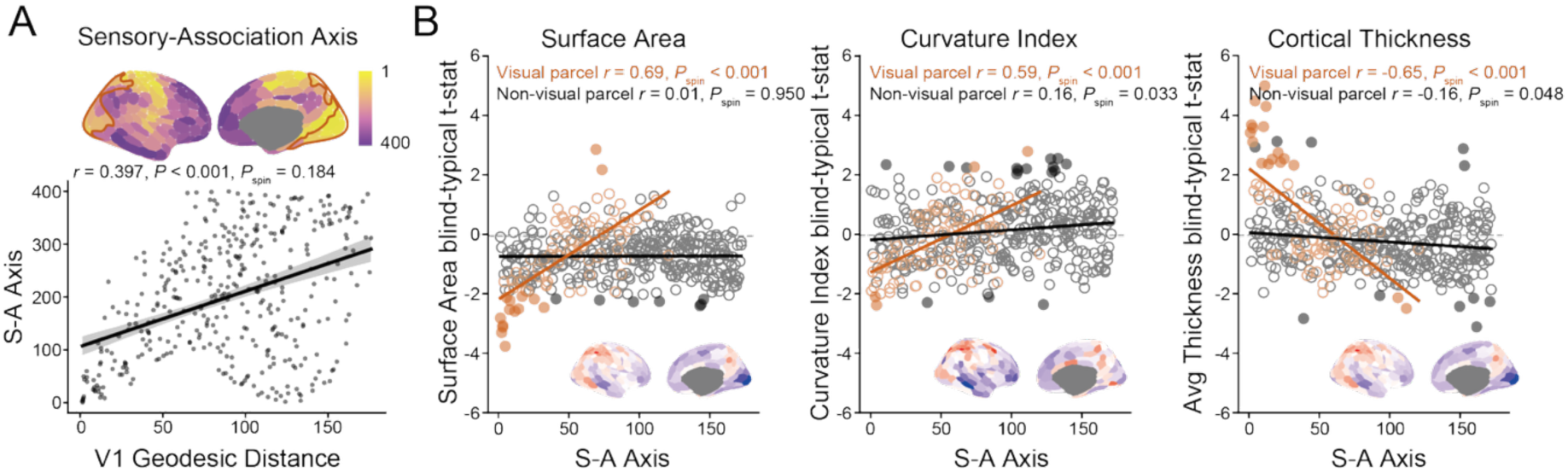
Cortical structural reorganization along sensorimotor-association axis in early blindness. **(A)** Correlation between geodesic distance from V1 and sensorimotor-association axis (*r*(398) = 0.397, *P* < 0.001, *P*spin = 0.184). **(B)** Blind–neurotypical t-statistics for surface area, curvature index, and cortical thickness across Schaefer 400 parcels, varied as a function of sensorimotor-association axis (surface area: *r*(84) = 0.69, *P* < 0.001, *P*spin < 0.001; curvature: *r*(84) = 0.59, *P* < 0.001, *P*spin < 0.001; thickness: *r*(84) = −0.65, *P* < 0.001, *P*spin < 0.001). Parcels overlapping visual regions are shown in orange, and non-overlapping parcels in black. Filled dots indicate statistical significance (*P* < 0.05). Surface maps in the bottom-right panels show the spatial distribution of blind–neurotypical *t*-statistics for each feature, standardized to a range of −2 to 2.

**Supplementary Figure 8.**
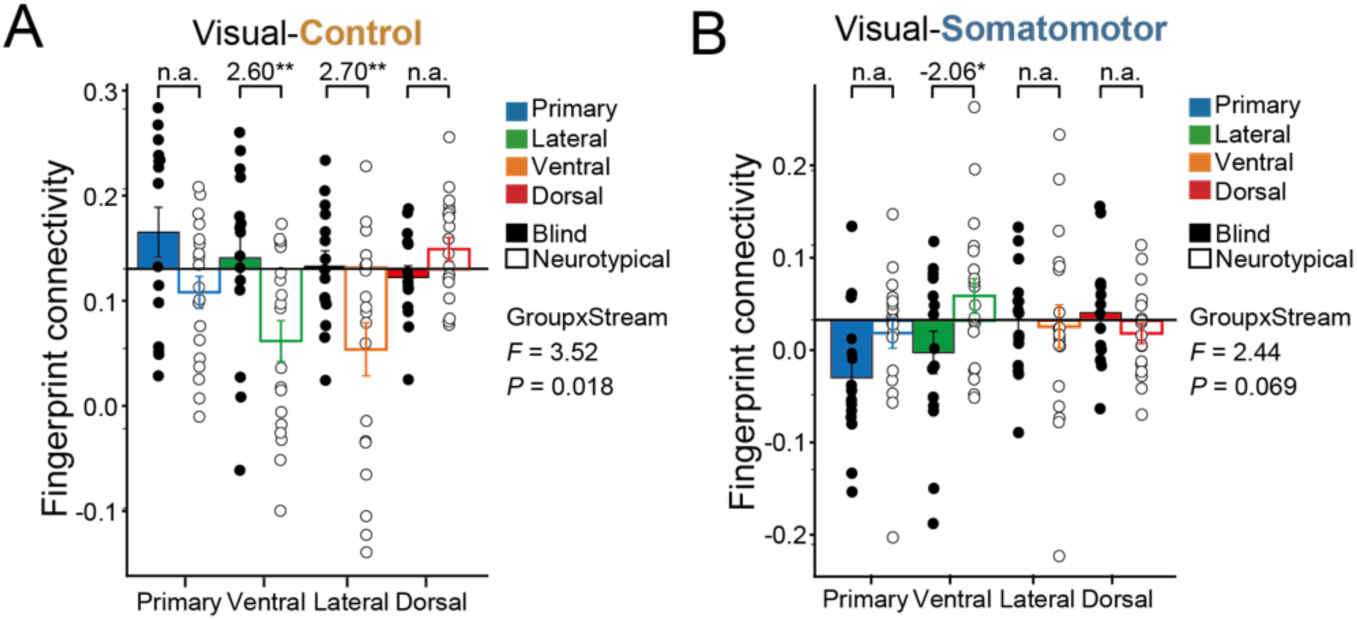
Cross-network thalamocortical plasticity differs across visual processing streams in early blindness. **(A)** Cortical visual–thalamic control functional fingerprint similarity in blind (filled) and neurotypical (open) participants, shown separately for primary, ventral, lateral, and dorsal visual processing streams. Visual–control fingerprint similarity showed a significant group × stream interaction, with smaller group differences in the dorsal pathway than in the ventral and lateral pathways. **(B)** Cortical visual–thalamic somatomotor functional fingerprint similarity in blind (filled) and neurotypical (open) participants, shown separately for primary, ventral, lateral, and dorsal visual processing streams.

**Supplementary Figure 9.**
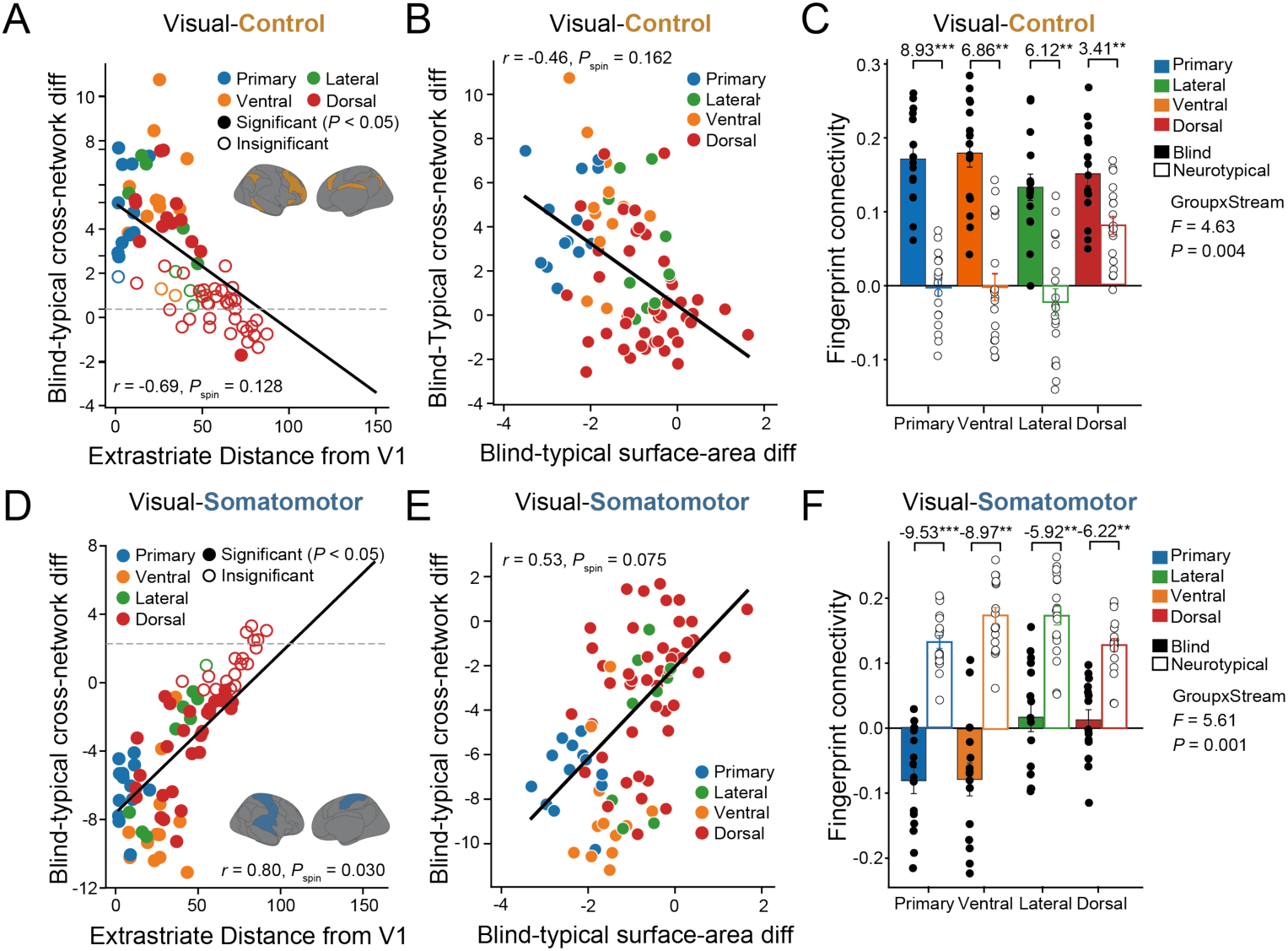
Hierarchical cortical functional reorganization in early blindness. **(A)** Blind–neurotypical tstatistics for visual–control functional fingerprints across the 86 Schaefer parcels overlapping visual regions, plotted as a function of geodesic distance from V1. Filled dots indicate statistical significance (*P* < 0.05). Colors denote processing streams (blue: primary, orange: ventral, green: lateral, red: dorsal). **(B)** Correlation between V1 surface area (blind vs. neurotypical t-values) and visual–control fingerprint similarity (blind vs. neurotypical t-values) across cortical parcels. **(C)** Visual–control network functional fingerprint similarity in blind (filled) and neurotypical (open) groups, shown separately for each processing stream. **(D)** Blind–neurotypical t-statistics for visual–somatomotor functional fingerprints across the same 86 parcels, plotted as a function of geodesic distance from V1. Filled dots indicate statistical significance (*P* < 0.05).**(E)** Correlation between V1 surface area (blind vs. neurotypical t-values) and visual–somatomotor fingerprint similarity (blind vs. neurotypical t-values) across cortical parcels. **(F)** Visual–somatomotor network functional fingerprint similarity in blind (filled) and neurotypical (open) groups, shown separately for each processing stream.

**Supplementary Figure 10.**
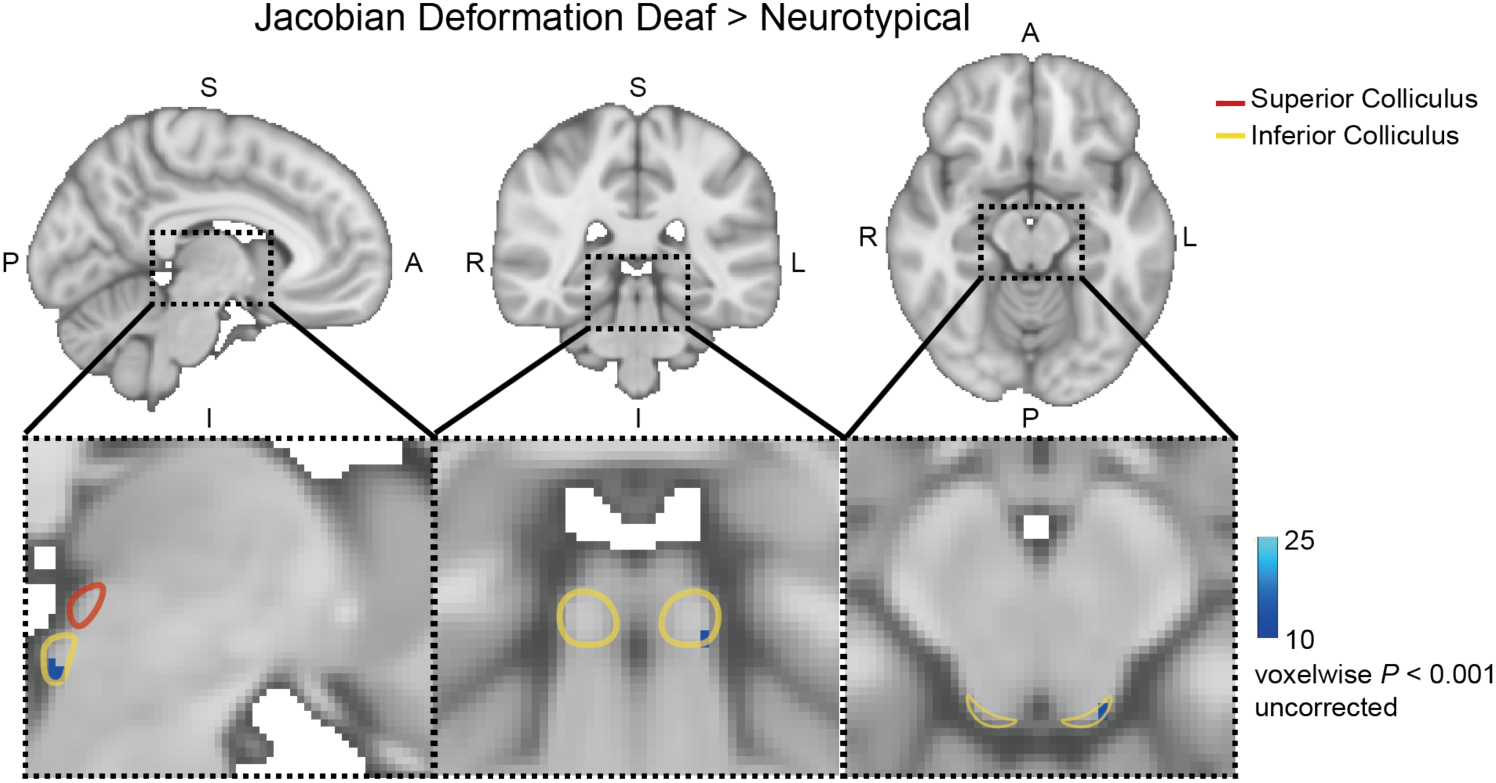
Structural differences in the inferior colliculus in early deafness. Voxels showing significantly lower log-Jacobian values in deaf compared with neurotypical participants are shown in blue (1,000 permutation tests, *P* < 0.001). Lower values indicate greater local compression during registration to the MNI template and therefore larger native anatomy relative to the template. The superior and inferior colliculi are outlined in red and yellow, respectively.

**Supplementary Figure 11.**
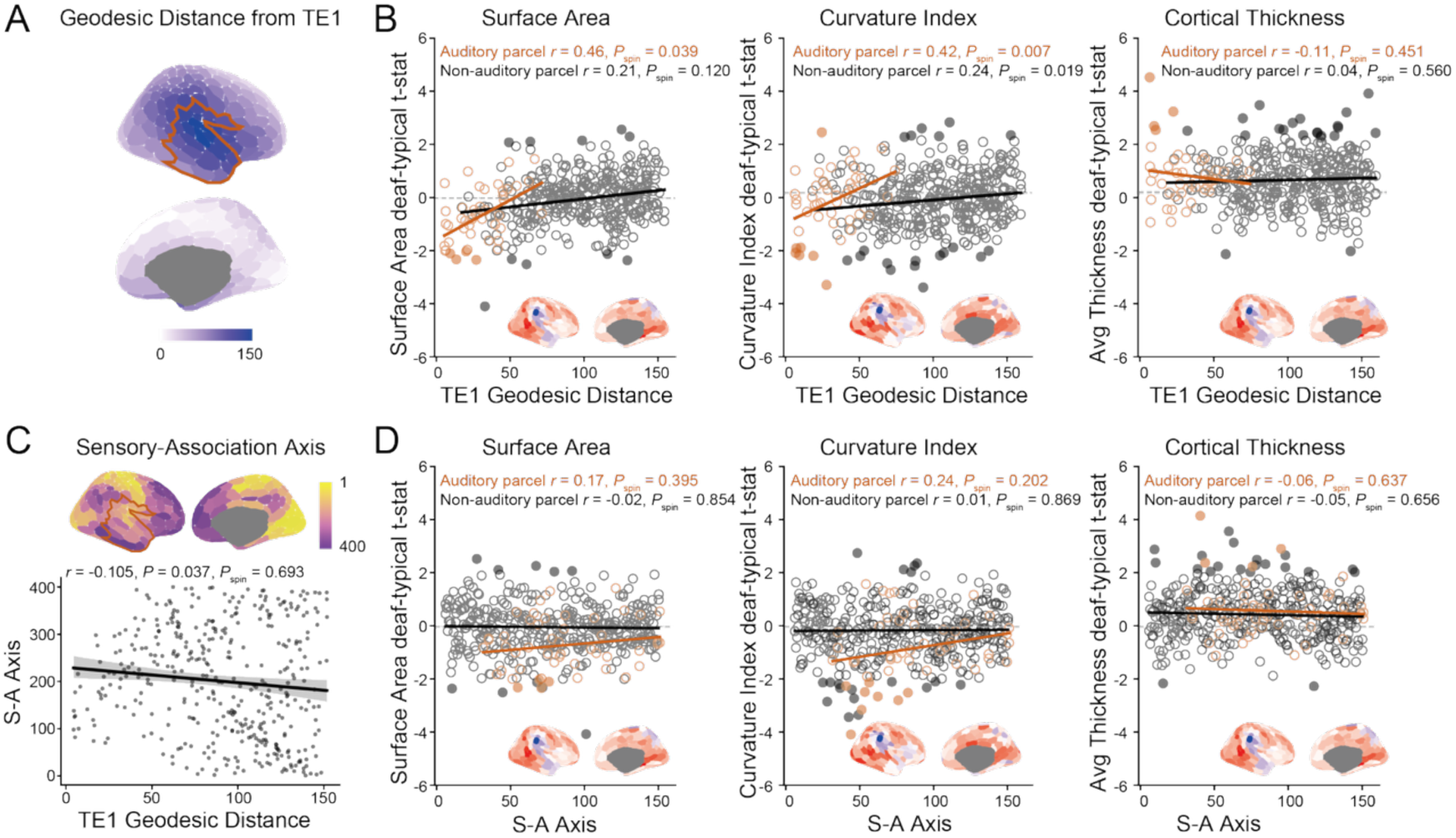
Cortical structural reorganization along hierarchy in early deafness. **(A)** Geodesic distance from TE1, overlaid with the auditory region ROI (orange). **(B)** Deaf–neurotypical t-statistics for surface area, curvature index, and cortical thickness across Schaefer 400 parcels, plotted as a function of geodesic distance from TE1. Parcels overlapping auditory regions are shown in orange, and non-overlapping parcels in black. Surface maps show the spatial distribution of each feature, standardized to a range of −2 to 2. **(C)** Correlation between geodesic distance from TE1 and sensorimotor-association axis (*r*(398) = −0.105, *P* = 0.037, *P*spin = 0.693). **(D)** Deaf–neurotypical t-statistics for surface area, curvature index, and cortical thickness across Schaefer 400 parcels showed no systematic relationship to sensorimotor-association axis (surface area *r*(47) = 0.17, *P* = 0.244, *P*spin = 0.395, curvature *r*(47) = 0.24, *P* = 0.097, *P*spin = 0.202; thickness *r*(47) = −0.062, *P* = 0.670, *P*spin = 0.637). Parcels overlapping auditory regions are shown in orange, and non-overlapping parcels in black. Filled dots indicate statistical significance (*P* < 0.05). Surface maps in the bottom-right panels show the spatial distribution of deaf–neurotypical *t*-statistics for each feature, standardized to a range of −2 to 2.

**Supplementary Figure 12.**
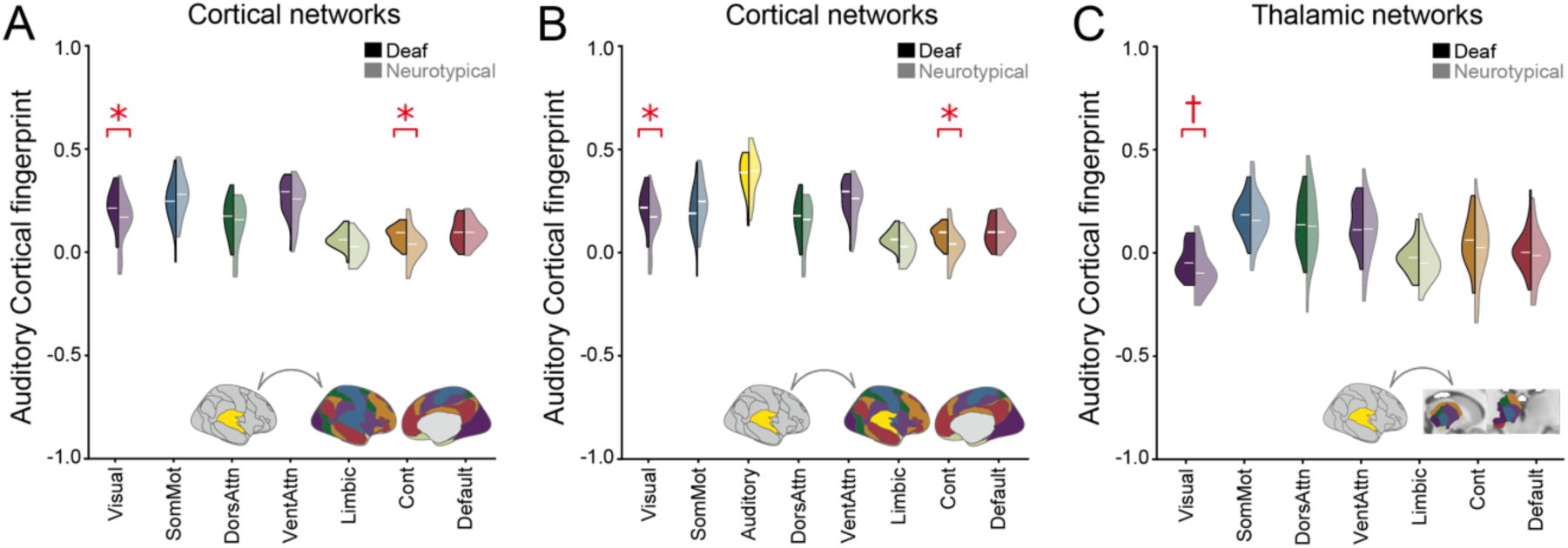
Cortical functional reorganization in early deafness. **(A, B)** Functional fingerprint similarity of the cortical auditory network (restricted to parcels overlapping the somatomotor network) with cortical Yeo networks, using either **(A)** the entire somatomotor network or **(B)** the somatomotor network subdivided into auditory-related and non-auditory subnetworks, in deaf and neurotypical groups. **(C)** Functional fingerprint similarity of the cortical auditory network (restricted to parcels overlapping the somatomotor network) with thalamic networks in deaf and neurotypical groups. \*\*\**P* < 0.001 (FDR-corrected), \*\**P* < 0.01 (FDR-corrected), \**P* < 0.05 (FDR-corrected). †*P* < 0.05 (FDR-uncorrected). FDR correction was applied across 7 (A, C) or 8 (B) networks.

**Supplementary Figure 13.**
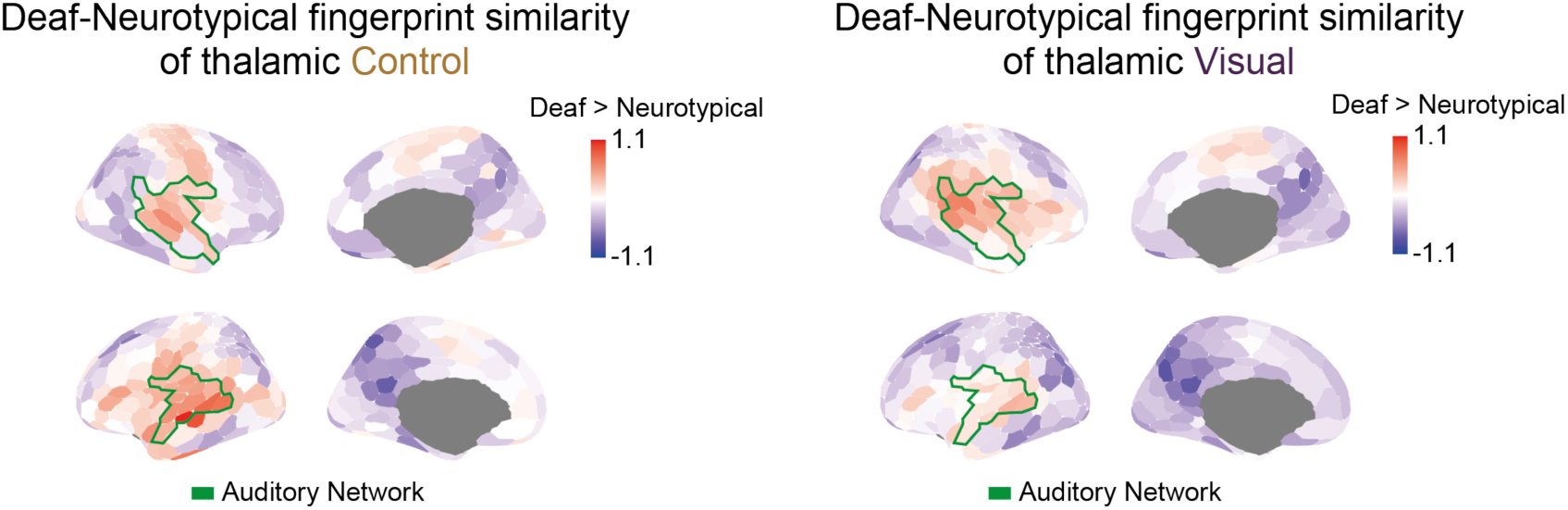
Thalamocortical functional reorganization in early deafness. Group differences between early deaf and neurotypical participants in the fingerprint similarity between the thalamic control and visual networks and cortical parcels. Cortical parcels overlapping with the auditory network are highlighted in green.

**Supplementary Figure 14.**
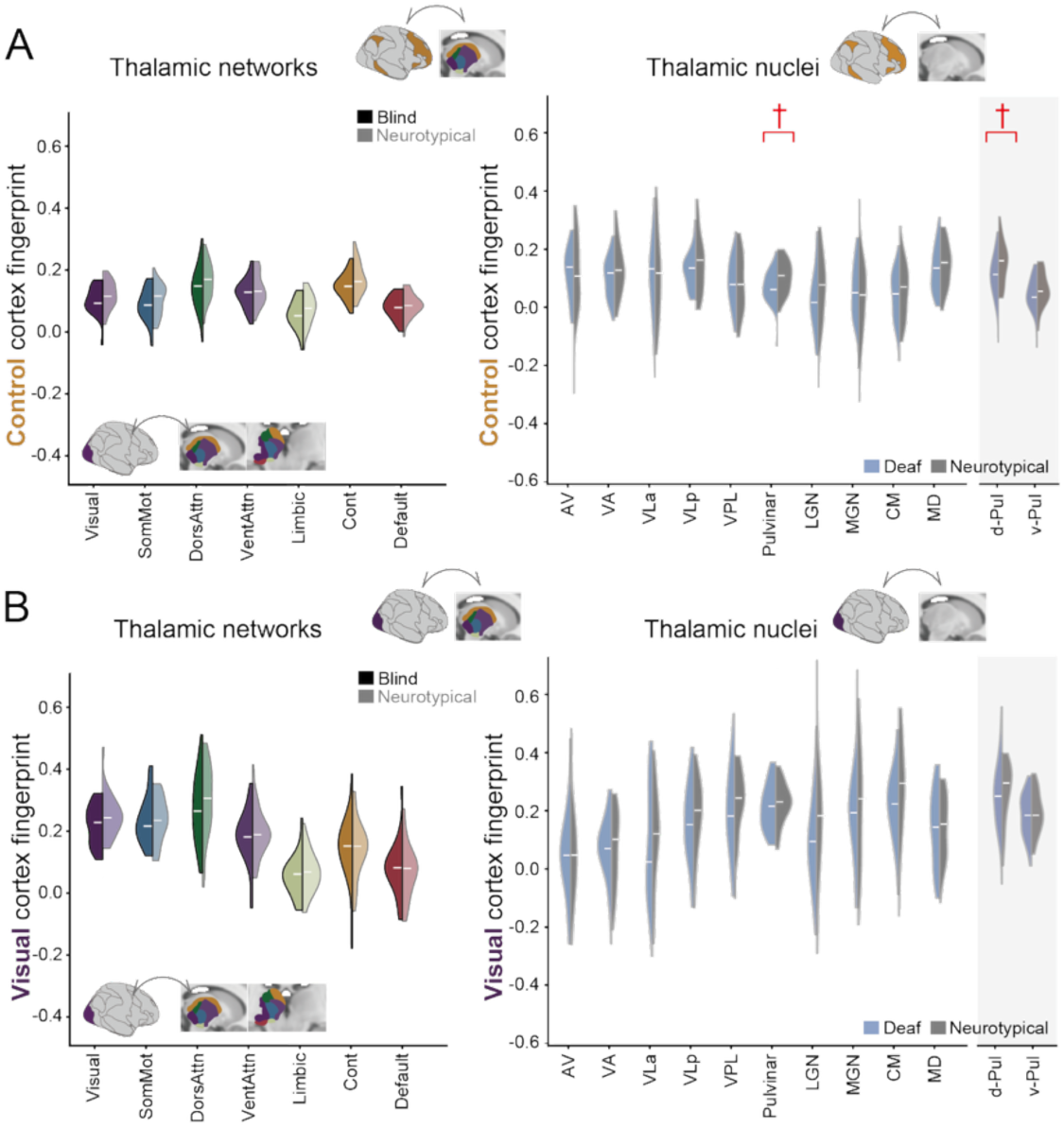
Thalamic functional reorganization in early deafness. **(A, B)** Functional fingerprint similarity of the cortical (A) control and (B) visual networks and thalamic nuclei in deafness. \**P* < 0.05 (FDR-corrected), †*P* < 0.05 (FDR-uncorrected). FDR correction was applied across 7 networks or 10 thalamic nuclei.

